# cpSRP54 and FtsH cooperate thylakoid membrane associated proteostasis during de-etiolation in *Arabidopsis thaliana*

**DOI:** 10.1101/2022.12.01.518768

**Authors:** Yang Lei, Bilang Li, Xiaomin Wang, Junyou Wei, Peiyi Wang, Jun Zhao, Fei Yu, Yafei Qi

**Author notes:** These authors contributed equally to this article (Yang Lei, Bilang Li). Corresponding author, Author for correspondence: Yafei Qi, State Key Laboratory of Crop Stress Biology for Arid Areas and College of Life Sciences, Northwest A&F University, Yangling, Shaanxi 712100, People’s Republic of China.

## Abstract

The thylakoid membrane protein quality control, which requires the coordination of membrane protein translocation and degradation of unassembled proteins, determines the chloroplast development during de-etiolation. Despite numerous efforts, the regulation of this progress in land plants is largely unknown. Here, we reported the isolation and characterization of *pga4* mutants with defects in chloroplast development during de-etiolation. Map-based cloning and complementation assay confirmed that *PGA4* encodes the chloroplast signal recognition particle 54 kDa protein (cpSRP54). A heterogeneous LhcB2-GFP was generated as an indicative substrate for cpSRP54-mediated thylakoid translocation. LhcB2-GFP were not assembled into functional complexes, and degraded to a short form dLhcB2-GFP during de-etiolation, through an N-terminal degradation initiated on thylakoid membranes. Further biochemical and genetic evidences demonstrated that the degradation of LhcB2-GFP to dLhcB2-GFP was disrupted in *pga4*, and *var2* mutants caused by mutations in the VAR2/AtFtsH2 subunit of thylakoid FtsH. The yeast two-hybrid assay showed that the N-terminus of LhcB2-GFP which was degraded consequently, interacts with the protease domain of VAR2/AtFtsH2 in yeasts. Moreover, the over-accumulated LhcB2-GFP in *pga4* and *var2*, formed protein aggregates, which were insoluble in mild nonionic detergents. Genetically, *cpSRP54* is a new suppressor locus for the leaf variegation phenotype of *var2*. Those together demonstrated the coordination of cpSRP54 and thylakoid FtsH in maintenance of thylakoid membrane protein quality control during the assembly of photosynthetic complexes, and provided a trackable substrate and product for monitoring the cpSRP54-dependent protein translocation and the FtsH-dependent protein degradation.

**One Sentence Summary:** We revealed the coordination of cpSRP54 and FtsH in thylakoid membrane protein quality control, and provided a trackable marker for monitoring the activity of cpSRP54 and thylakoid FtsH protease.

## Introduction

During chloroplast development, the proper biogenesis of photosynthetic protein complexes requires translocation of apoproteins into the membrane, assembly of each subunit with a right stoichiometry, as well as the tight regulation of biosynthesis and incorporation of photosynthetic pigments (Pribil et al., 2014; Wang and Grimm, 2015; Zhu et al., 2022). Chloroplast signal recognition particle 54 kDa protein (cpSRP54) is the plant homolog of bacterial SRP54, involved in co-translational targeting of nascent chloroplast proteins to thylakoid membrane (Akopian et al., 2013; Hristou et al., 2019). In addition, cpSRP54 is also required for translocation of light-harvesting chlorophyll a/b binding proteins (LHCP) to thylakoids, by forming a transit complex with cpSRP43 and the hydrophobic client LHCP (Schünemann et al., 1998; Yuan et al., 2002; Goforth et al., 2004; Holdermann et al., 2012; Ziehe et al., 2018). The assembly of LHCP on thylakoid membranes is tightly coordinated with the biosynthesis of chlorophylls (Wang and Grimm, 2015; Wang et al., 2018). For instance, in Arabidopsis *chlorina* mutants lacking chlorophyll *b*, caused by the lesion of chlorophyll *a* oxygenase (CAO), the accumulation of LHC apoprotein was reduced drastically, mostly likely due to a proteolytic degradation performed by yet unknown chloroplast proteases (Reinbothe et al., 2006). These indicated that the chloroplast PQC system could work cooperatively with the membrane protein targeting or assembly system, when highly hydrophobic photosynthetic apoproteins are prone to be aggregated, misfolded or photo-damaged during the assembly of photosynthetic complex. Nonetheless, a convenient and trackable substrate protein marker to dissect the coordination work *in vivo*, has not been reported in land plants.

FtsH (*filamentous temperature sensitive H*), belonged to the AAA (<u>A</u>TPases <u>a</u>ssociated with a variety of cellular <u>a</u>ctivities) protease family, is a specific type of membrane-localized metalloprotease to maintain membrane protein quality control (PQC). FtsH exists universally in almost all prokaryotic species, and organelles of eukaryotic cells such as mitochondria and chloroplasts (Ogura et al., 1991; Tomoyasu et al., 1995; Arlt et al., 1996; Sakamoto et al., 2003). In *E. coli*, FtsH is responsible for the degradation of numerous substrates with various physiological functions (Ito and Akiyama, 2005; Langklotz et al., 2012) including heat shock sigma factor σ32 (Herman et al., 1995; Tomoyasu et al., 1995). In yeasts, two types of FtsH homologs are anchored to the inner membrane of mitochondria, including the i-AAA protease YME1 facing to inter membrane space and the i-AAA protease YTA10/12 to the matrix side (Arlt et al., 1996; Weber et al., 1996; Glynn, 2017). The proposed FtsH working model for membrane PQC is that the ATPase domain performs as an unfoldase to unfold its substrates to polypeptide chains, which are subsequently translocated to the proteolytic domain for degradation (Suno et al., 2006; Bieniossek et al., 2009; Puchades et al., 2017).

In *Arabidopsis*, thylakoid FtsH complex are heterohexamers, composed of type A (AtFtsH1 and 5) and type B (AtFtsH2 and 8) subunits, with the ATPase domain and the protease domain facing to the stroma side (Sakamoto et al., 2003; Yu et al., 2004; Zaltsman et al., 2005). Mutations in *VAR2/AtFtsH2* and *VAR1/AtFtsH5* caused pronounced phenotype of variegated rosette leaves in *yellow variegated2* (*var2*) and *var1* respectively (Chen et al., 2000; Sakamoto et al., 2002). Complete loss of type A or type B leads to albino seedlings, indicating that thylakoid FtsH is essential for chloroplast development (Yu et al., 2005; Zaltsman et al., 2005). Taking advantage of the leaf variegation phenotype of *var2*, a set of genetic modifier mutants and their loci, including *Suppressors of variegation* (*SVRs*) and *Enhancers of variegation* (*EVRs*), were identified recently (Zheng et al., 2016; Wang et al., 2018; Liu et al., 2019; Qi et al., 2020; Wang et al., 2022). One group of those suppressor loci encode proteins that participate in chloroplast translation directly, including the chloroplast ribosomal protein RPL24 (Liu et al., 2013), the chloroplast translation initiation factor IF3 (Zheng et al., 2016), and the chloroplast translation elongation factor EF-Tu (Liu et al., 2019).These SVRs and EVRs demonstrated that the balance of chloroplast translation and cytosol translation regulates chloroplast development and leaf variegations (Wang et al., 2018).

Similar to mitochondrial inner membrane for oxidative phosphorylation (OXPHOS), thylakoid membrane of chloroplast is where photophosphorylation takes place. Driven by light energy, the electron transfer reaction is achieved by photosynthetic protein complexes that embedded in thylakoid membranes, including photosystem II (PSII), cytochrome *b_6_f* (Cyt*b_6_f* and photosystem I (PSI). Among them, PSII is the most vulnerable, and its reaction center proteins are prone to be photo-damaged under high light stress (Aro et al., 1993; Nixon et al., 2010). FtsH plays essential roles in the protein quality control of photosynthetic protein complexes, for instance, by degrading photo-damaged PSII reaction center protein D1 (Lindahl et al., 2000; Bailey et al., 2002; Silva et al., 2003; Malnoё et al., 2014). Under other abiotic stress conditions such as sulfur deprivation or dark induced senescence, other potential substrates of thylakoid FtsH had also been reported, including Cyt*b_6_f* and light-harvesting complex LHCII and LHCI (Luciński and Jackowski, 2013; Malnoё et al., 2014; Bujaldon et al., 2017). However, the assessment of D1 fragment degradation is difficult under nonphotoinhibitory light conditions in *Arabidopsis* (Kato et al., 2012).

The protein processing activity of FtsH requires the presence of a stable domain existing in the substrate, as the unfoldase activity of FtsH is not robust, and it will release the processed substrate when encountering the stable domain (Okuno et al., 2006; Okuno and Ogura, 2013). In this work, we generated a transgenic line expressing a chimeric LhcB2-GFP to dissect the regulation of chloroplast PQC and the processing activity of FtsH. Using *pale green Arabidopsis* mutants, *pga4-1* and *pga4-2* with disruptions in *cpSRP54*, we confirmed that the chimeric LhcB2-GFP was targeted to thylakoid membrane by cpSRP54, and degraded to dLhcB2-GFP mediated by thylakoid FtsH through an N-terminal processing. More interestingly, *PGA4/cpSRP54* is a new genetic suppressor locus of *var2* leaf variegation. Our work presented an indicative marker to dissect the proteolytic activity of thylakoid FtsH, and most importantly, provided a new insight that cpSRP54 and thylakoid FtsH function together for chloroplast PQC during de-etiolation.

## Results

### *PGA4* encodes cpSRP54

To study the mechanisms underlying chloroplast protein homeostasis, we systematically isolated and characterized *pale green Arabidopsis* (*pga*) mutants from our ethyl methanesulfonate (EMS) mutagenesis pools (Li et al., 2022). The *pga4-1* showed pale green leaves, indicating defects in chloroplast development and accumulation of photosynthetic pigments and protein complex (Supplemental Figure 1A). We employed the map-based cloning approach to identify the *PGA4* locus (Figure 1A). Initial mapping placed *PGA4* on chromosome 5, and further fine mapping narrowed down the location of *PGA4* near to the marker *F8F6#1* (Figure 1A). DNA sequencing identified a G to A mutation in the coding region of AT5G03940, resulting a nonsense mutation (TGA) in the 89^th^ amino acid residue tryptophan (TGG) of cpSRP54 (Figure 1A). In addition, the second allele of *PGA4* mutant, *pga4-2* recovered from an activation tagging mutagenesis pool (Yu et al., 2008), was confirmed by the allelism test with *pga4-1* and PCR-based genotyping (Supplemental Figure 1A-C). qRT-PCR confirmed that the accumulation of *cpSRP54* transcripts is severely decreased in *pga4-2* (Supplemental Figure 1D).

**Figure 1.**
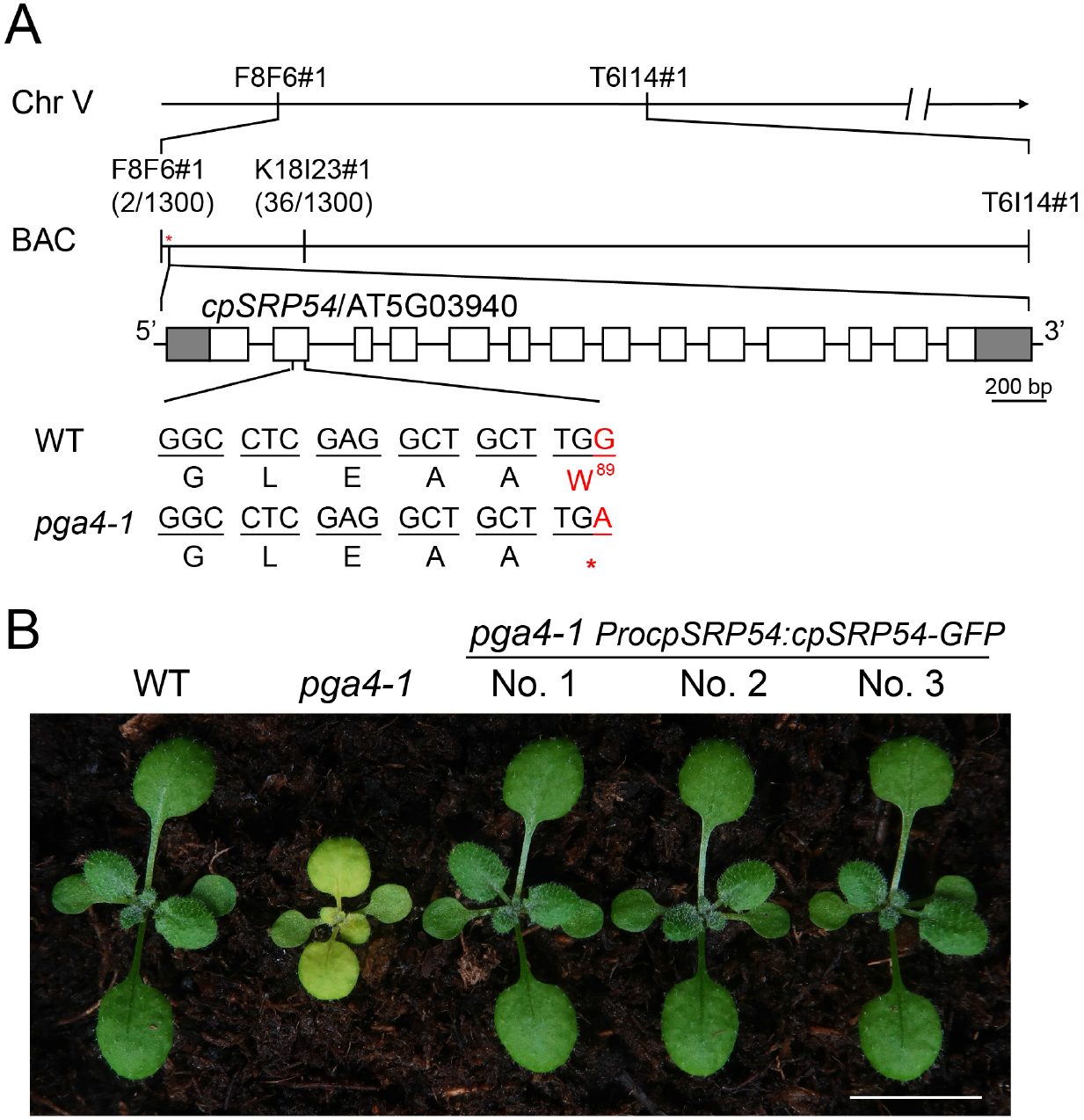
*PGA4* encodes cpSRP54. A. The Map-based cloning of *pga4-1*. The *PGA4* locus was mapped between *F8F6#1* and *K18I23#1* using 650 individual plants. In this region, a nonsense mutation (Trp/W^89^ to a stop codon) in *cpSRP54/ AT5G03940* was identified. Gray and white boxes represent UTRs and exons of *cpSRP54/*AT5G03940, respectively. B. Complementation of *pga4-1* with the binary construct *ProcpSRP54:cpSRP54-GFP*.

For complementation assay, a plant binary vector *ProcpSRP54:cpSRP54-GFP* was constructed, in which the genomic DNA of *cpSRP54* is driven by its native promoter and fused with coding sequences of green fluorescent protein (GFP) at its 3’ end. The pale-green *pga4-1* was recovered to the wild-type phenotype using this vector (Figure 1B). In *pga4-1 ProcpSRP54:cpSRP54-GFP* transgenic lines, cpSRP54-GFP were accumulated in etioplasts (0 h) and chloroplasts (24 h) during de-etiolation (Supplemental Figure 2). These finally confirmed that the pale-green phenotype of *pga4-1* and *pga4-2* were caused by mutations in *cpSRP54*.

### Generation of a stable transgenic line expressing a chimeric LhcB2-GFP

As cpSRP54 is required for targeting LHC proteins to thylakoid membranes, we tried to express a heterogeneous LhcB2-GFP as a client for cpSRP54. We constructed a binary vector *ProLhcB2:gLhcB2-GFP*, in which the genomic DNA of AT2G05070 (*LhcB2.2*) was driven by its native promoter and fused with the GFP coding sequence at its 3’ end (Figure 2A). To confirm the expression of *LhcB2-GFP* is consistent with the endogenous *LhcB2*, we firstly transformed this construct into the wild-type Arabidopsis (WT). One representative WT *ProLhcB2:gLhcB2-GFP* line (hereafter designated WT *LhcB2-GFP*) was treated with lincomycin (Lin) and norflurazon (NF) that could block chloroplast translation and carotenoid biosynthesis respectively (Susek et al., 1993; Mulo et al., 2003). Chemical treatments led to albino seedlings as expected (Figure 2B). qRT-PCR analyses revealed that the accumulation of *LhcB2-GFP* and *LhcB2* transcripts were severely reduced in Lin or NF treated seedlings compared with those in the mock (Figure 2C). Those together indicated that the expression of *LhcB2-GFP* in WT *LhcB2-GFP* is regulated by retrograde repression, being consistent with the endogenous *LhcB2*.

**Figure 2.**
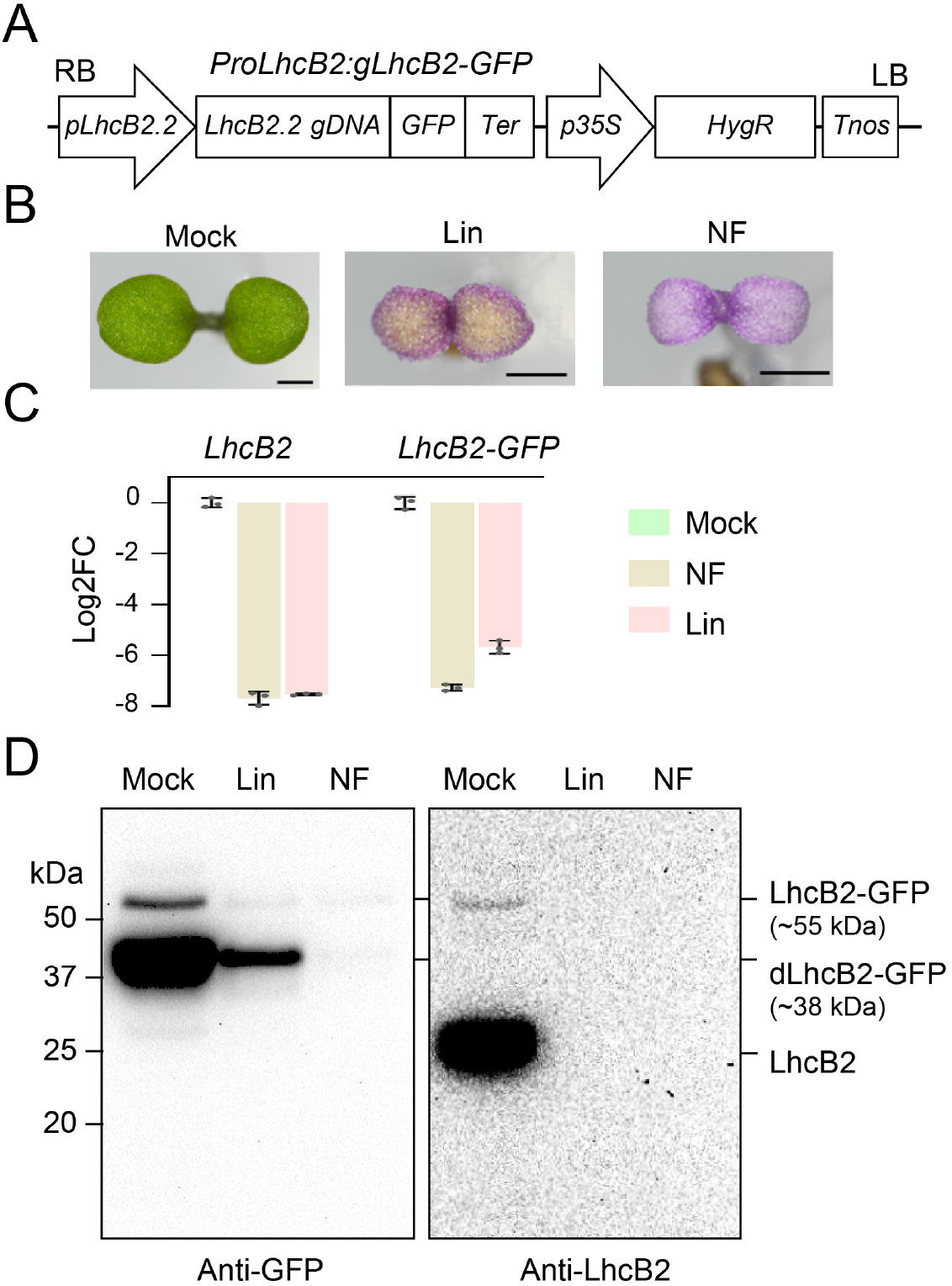
The generation of a transgenic line WT *LhcB2-GFP*. A. The vector construct of *ProLhcB2:gLhcB2-GFP*. B. Seedlings of WT *LhcB2-GFP* were grown on 1/2 MS (Mock), and 1/2 MS containing 0.5 mM lincomycin (Lin) or 5 μM norflurazon (NF) for 4 days. Bars, 1 mm. C. qRT-PCR analyses of accumulations of *LhcB2* and *LhcB2-GFP* in WT *LhcB2-GFP* treated with lincomycin and norflurazon as in (B). Base 2 logarithm of the fold-change (Log2FC) were calculated, and *PP2A* was used as the reference gene. Data are means ± s.d. of three biological replicates. D. The accumulation of LhcB2-GFP (~55 kDa) and dLhcB2-GFP (~38 kDa) in 4-day-old WT *LhcB2-GFP* treated with Lin and NF as in (B). An anti-GFP antibody and an anti-LhcB2 antibody were used, respectively. Protein loading was normalized to equal fresh tissue weight.

The accumulation of LhcB2-GFP in those seedlings was examined with immunoblotting, using an anti-GFP antibody and an anti-LhcB2 antibody. A full-length LhcB2-GFP (~55 kDa), and a degradation form of LhcB2-GFP designated dLhcB2-GFP (~38 kDa), were highly accumulated in the mock compared with those in seedlings treated with Lin or NF (Figure 2D). The dLhcB2-GFP was only recognized by the anti-GFP antibody, but not by the anti-LhcB2 antibody (Figure 2D). Consistently, reduced GFP and chlorophyll fluorescence were observed in cotyledon chloroplasts in seedlings treated with Lin or NF, compared with the mock (Supplemental Figure 3).

We also examined the accumulation of LhcB2-GFP during 24-h de-etiolation (Supplemental Figure 4). This assay could provide a rational starting point to analyze the accumulation of photosynthetic proteins (Qi et al., 2020; Pipitone et al., 2021). Consistent with those in 4-day-old seedlings, both LhcB2-GFP (~55 kDa) and dLhcB2-GFP (~38 kDa) were strongly accumulated in 24-h de-etiolated WT *LhcB2-GFP* (Figure 2D and Supplemental Figure 4A). Besides, a larger form of signal band (~65 kDa) was observed in a longer exposure time, which was likely the LhcB2-GFP precursor (pLhcB2-GFP) containing the chloroplast transit peptide (cTP). As *LhcB2-GFP* was driven by its native promoter, the accumulation of LhcB2-GFP and dLhcB2-GFP were responsive to light induction (Supplemental Figure 4A). In 24-h de-etiolated WT *LhcB2-GFP*, the overlap of GFP and chlorophyll fluorescence confirmed that LhcB2-GFP is targeted into chloroplasts (Supplemental Figure 4B).

### The chimeric LhcB2-GFP could be target to thylakoid membranes during de-etiolation

The detection of full-length LhcB2-GFP enables us to check its chloroplastic localization, via chloroplast fractionation using 24-h de-etiolated seedlings of WT *LhcB2-GFP* (Figure 3A). Crude membrane fractions (P) and soluble fractions (S) were isolated, using a hypotonic buffer (HM) or HM buffer containing higher concentration of ionic salts (2 M NaCl). The large subunit of RuBisCO (RbcL, a stroma protein marker) was enriched in soluble fractions, while the endogenous LhcB2 (a thylakoid membrane protein marker) existed in the crude membrane fractions (Figure 3A). Partial LhcB2-GFP and dLhcB2-GFP were present in the S fraction when HM buffer were used, indicating that they were distributed in the stroma (Figure 3A). There were still some LhcB2-GFP and dLhcB2-GFP in the P fractions, and washing with 2 M NaCl buffer did not increase their proportion into the S fractions, suggesting that partial LhcB2-GFP and dLhcB2-GFP were either associated with the membrane fractions, or aggregated (Figure 3A).

**Figure 3.**
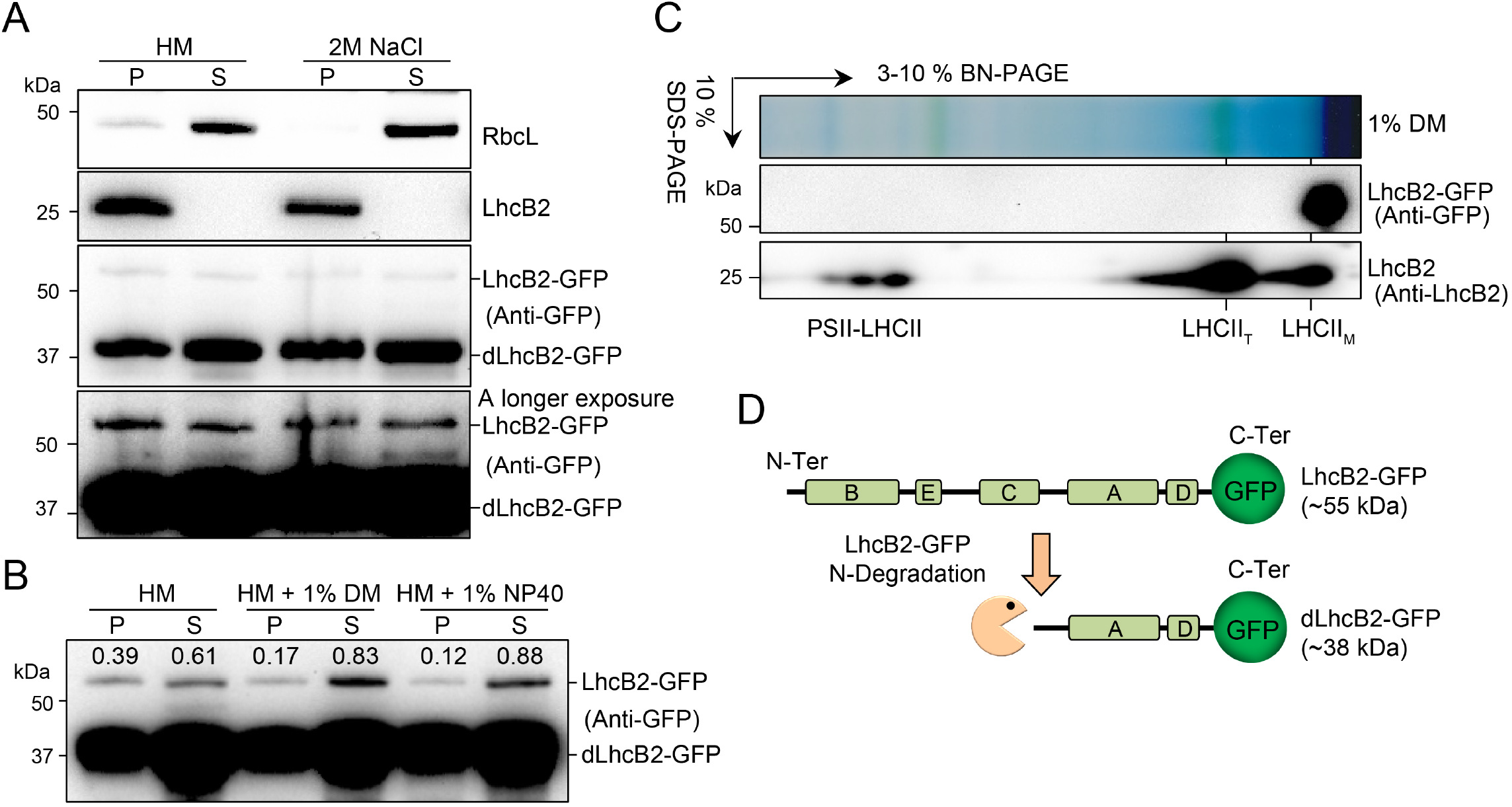
The heterogeneous LhcB2-GFP is not assembled into antenna complexes. A. Subcellular fractionation of LhcB2-GFP and dLhcB2-GFP in 24-h de-etiolated WT *LhcB2-GFP*. The soluble stroma fraction (S) was indicated by RbcL, and the insoluble fraction (P) by endogenous LhcB2. B. The membrane associated LhcB2-GFP were solubilized with two mild nonionic detergents n-dodecyl-β-D-maltoside (1% β-DM) and Nonidet P-40 (1% NP40). The accumulation of LhcB2-GFP in the solubilized and pellet fractions was relatively quantified with signal intensities. C. Thylakoid membrane of 24-h de-etiolated WT *LhcB2-GFP* were solubilized with 1% DM and resolved on 1-D BN-PAGE and the 2-D SDS-PAGE. The accumulation of LhcB2-GFP and LHCII complexes were detected with the anti-GFP antibody and the anti-LhcB2 antibody respectively. D. The diagram of the N-terminal degradation of LhcB2-GFP.

In order to check that if LhcB2-GFP and dLhcB2-GFP were associated with thylakoid membranes, we used two mild nonionic detergents, n-dodecyl-β-D-maltoside (β-DM) and Nonidet P-40 (NP40), to solubilize membrane proteins. Interestingly, in contrast to dLhcB2-GFP, more than half amounts of LhcB2-GFP in the P fraction (0.22 of 0.39 by β-DM, and 0.27 of 0.39 by NP40) were solubilized into the detergent-dissolved S fractions (Figure 3B). This suggested that LhcB2-GFP could be associated with thylakoid membranes, and dLhcB2-GFP was mainly in the stroma. However, there were still partial LhcB2-GFP and dLhcB2-GFP that were resistant to 1% β-DM or 1% NP40 (Figure 3B). These demonstrated that LhcB2-GFP were targeted onto thylakoid membranes, and detergent-resistant LhcB2-GFP and dLhcB2-GFP may form protein aggregates.

### The chimeric LhcB2-GFP is unassembled and degraded to dLhcB2-GFP through N-terminal degradation

The high level of dLhcB2-GFP suggested that LhcB2-GFP is not functional properly as endogenous LhcB2. We investigated the assembly of membrane-associated LhcB2-GFP and endogenous LhcB2 complexes (Figure 3C). Solubilized membrane proteins of 24-h de-etiolated WT *LhcB2-GFP* were resolved on 1-D BN-PAGE and followed by 2-D SDS PAGE, and detected with the anti-GFP antibody and the anti-LhcB2 antibody respectively. The endogenous LhcB2 forms multiple protein complexes as reported (Järvi et al., 2011), including LHCII monomer (LHCII_M_), LHCII trimer (LHCII_T_) and PSII-LHCII supercomplexes (PSII-LHCII) (Figure 3C). However, the membrane-associated LhcB2-GFP were not assembled into large functional LHCII antenna protein complexes, and present as a monomer (Figure 3C).

Based on the reported structure of LHC in *Spinacia oleracea*, LhcB2 proteins harbor five helices A to E, among which helices B, C, and A are transmembrane helices, spanning across thylakoid membrane (Figure 3D) (Liu et al., 2004). As the anti-LhcB2 antibody could detect endogenous LhcB2 and LhcB2-GFP, but not dLhcB2-GFP (Figure 2D), it was deduced that the anti-LhcB2 antibody may recognize the N-terminal region of LhcB2. To prove that, we expressed the recombinant LhcB2 proteins, including GST-LhcB2, GST-ΔN-LhcB2, and GST-LhcB2-ΔC in *Escherichia coli*. The partial N or C terminal segments of recombinant GST-ΔN-LhcB2 and GST-LhcB2-ΔC were removed respectively. GST-LhcB2, GST-ΔN-LhcB2, and GST-LhcB2-ΔC were well detected by an anti-GST antibody, whereas only GST-ΔN-LhcB2 was undetectable by the LhcB2 antibody (data not shown). Thus an N-terminal degradation of LhcB2-GFP was proposed, and chloroplast proteases are most likely involved in that (Figure 3D).

### Loss of cpSRP54 affects the translocation of LhcB2-GFP to thylakoid membranes and the degradation of LhcB2-GFP to dLhcB2-GFP

To check if the chimeric LhcB2-GFP is targeted to thylakoid by cpSRP54, we crossed *pga4* mutants with WT *LhcB2-GFP* to generate *pga4 LhcB2-GFPs*. During 24-h de-etiolation, *pga4 LhcB2-GFPs* showed almost completely delayed greening phenotypes and reduced accumulation of photosynthetic pigments, compared with those in WT *LhcB2-GFP* (Figure 4A and B). These indicated that cpSRP54 is required for chloroplast development during de-etiolation. The accumulation of LhcB2-GFP and dLhcB2-GFP were analyzed in 24-h de-etiolated WT *LhcB2-GFP* and *pga4-1 LhcB2-GFP* (Figure 4C). Strikingly, LhcB2-GFP was largely accumulated in de-etiolated *pga4-1 LhcB2-GFP* compared with those in de-etiolated WT *LhcB2-GFP* (Figure 4C). On the other hand, the level of dLhcB2-GFP was sharply reduced in de-etiolated *pga4-1 LhcB2-GFP* compared with that in de-etiolated WT *LhcB2-GFP* (Figure 4C). Interestingly, the accumulation of LhcB2 in *pga4-1 LhcB2-GFP* was comparable to that in WT *LhcB2-GFP* (Figure 4C). Similar observations on the accumulation of LHCPs had been reported in *cpsrp54/ffc* mutants (Amin et al., 1999). As chlorophylls were severely reduced, it was suggested that the accumulated LhcB2 are apoproteins in *pga4-1 LhcB2-GFP*.

**Figure 4.**
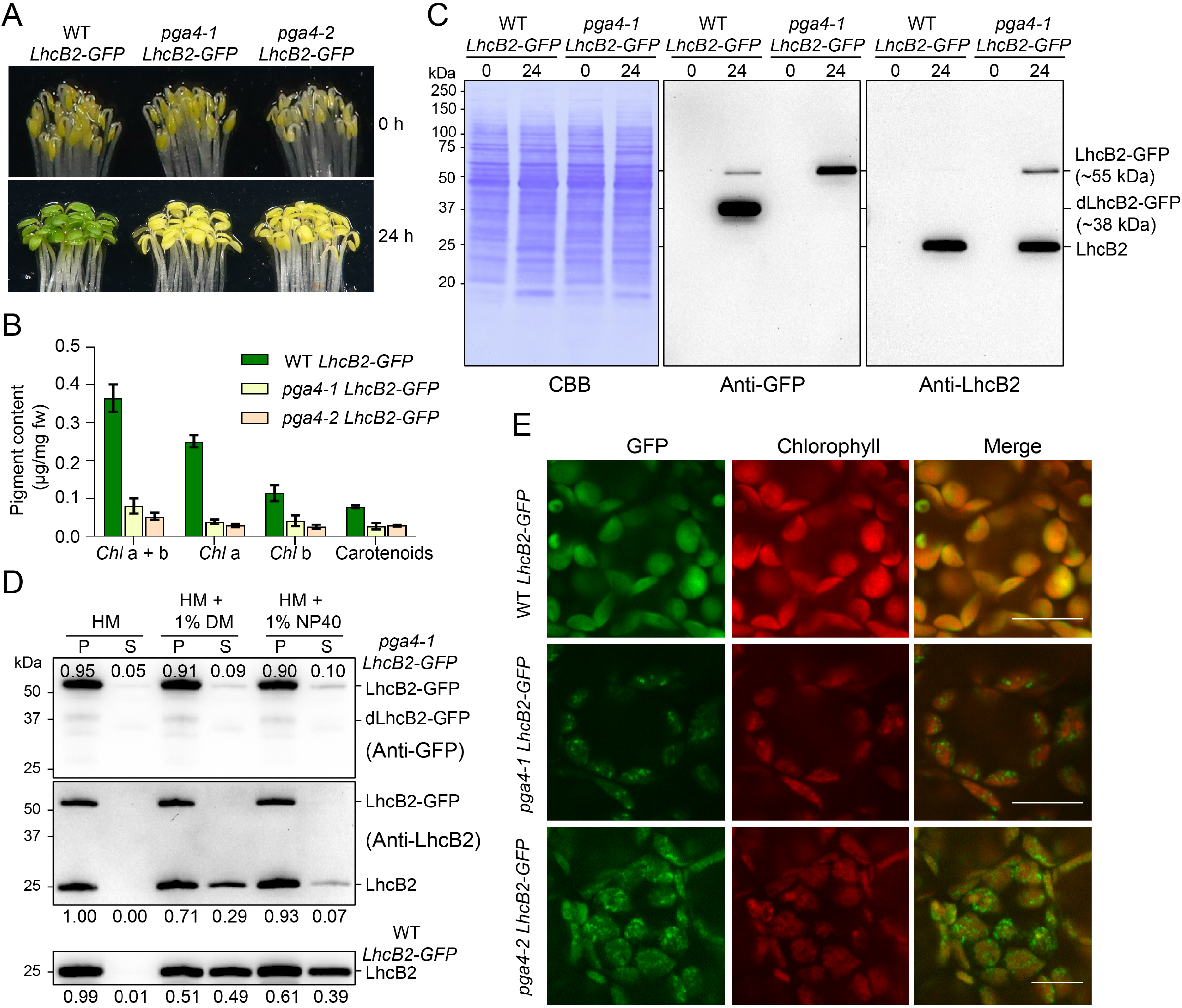
Loss of cpSRP54 affects the translocation of LhcB2-GFP to thylakoid membranes. A. Phenotypes of etiolated and 24-h de-etiolated WT *LhcB2-GFP, pga4-1 LhcB2-GFP*, and *pga4-2 LhcB2-GFP*. B. The accumulation of photosynthetic pigments in 24-h de-etiolated seedlings shown in (A). C. The accumulation of LhcB2-GFP and dLhcB2-GFP in 24-h de-etiolated WT *LhcB2-GFP* and *pga4-1 LhcB2-GFP*. Protein loading was normalized to equal fresh tissue weight. D. The solubility of LhcB2-GFP and LhcB2 in 24-h de-etiolated *pga4-1 LhcB2-GFP*. Mild nonionic detergents 1% β-DM and 1% NP40 were used to solubilize membrane proteins. The anti-GFP antibody and the anti-LhcB2 antibody were used, respectively. The accumulation of LhcB2-GFP and LhcB2 in the P and S fractions was relatively quantified with signal intensities. E. The detection of GFP and chlorophyll fluorescence in 24-h de-etiolated WT *LhcB2-GFP*, *pga4-1 LhcB2-GFP*, and *pga4-2 LhcB2-GFP*. DIC for differential interference contrast images. Bars, 10 μm.

We next examined the solubility of LhcB2-GFP and LhcB2 in 24-h de-etiolated *pga4-1 LhcB2-GFP*. Surprisingly, most of LhcB2-GFP (about 90%) may present as protein aggregates in the P fraction after 1% β-DM or 1% NP40 treatment in de-etiolated *pga4-1 LhcB2-GFP* (Figure 4D). Similarly, a higher relative level of detergent-resistant LhcB2 (71% with 1% β-DM, 93% with 1% NP40) were accumulated in *pga4-1 LhcB2-GFP* (Figure 4D), in comparison with those (51% with 1% β-DM, 61% with 1% NP40) in WT *LhcB2-GFP*. The detergent-dissolved LhcB2 in WT *LhcB2-GFP* is higher than those in *pga4-1 LhcB2-GFP* (Figure 4D). These indicated that both LhcB2-GFP and endogenous LhcB2 were clients of cpSRP54 for thylakoid membrane targeting, and cpSRP54 prevents their aggregations.

As LhcB2-GFP was highly detected in *pga4-1 LhcB2-GFP*, we next checked the localization of LhcB2-GFP in chloroplasts. The GFP fluorescence in 24-h de-etiolated seedlings was monitored with a confocal microscope. Surprisingly, it was shown that LhcB2-GFP formed discrete foci, surrounding chlorophyll fluorescence in 24-h de-etiolated *pga4-1 LhcB2-GFP* and *pga4-2 LhcB2-GFP* (Figure 4E). These discrete foci supported that over-accumulated LhcB2-GFP formed protein aggregates in *pga4 LhcB2-GFPs in vivo*, and being resistant to mild nonionic detergents. These together indicated that cpSRP54 is involved in targeting the chimeric LhcB2-GFP to thylakoids, where potential proteases for the N-terminal degradation of LhcB2-GFP to dLhcB2-GFP are localized.

### Thylakoid FtsH protease is required for LhcB2-GFP degradation

Thylakoid FtsH is the most abundant AAA protease on thylakoid membranes. Reduced activity of thylakoid FtsH caused by mutations in *VAR2/AtFtsH2*, led to *yellow variegated2* (*var2*) alleles with different extents of variegated rosette leaves. *var2-5* is a weaker allele in leaf variegation compared with *var2-4* (Chen et al., 2000; Yu et al., 2004). To find out if thylakoid FtsH is involved in the degradation of LhcB2-GFP, we obtained *var2-4 LhcB2-GFP* and *var2-5 LhcB2-GFP* via genetic crosses. In contrast to the weaker allele *var2-5*, *var2-4 LhcB2-GFP* showed a visible pale-green phenotype at 24-h de-etiolation, similar to those observed in *pga4 LhcB2-GFPs* (Figure 5A and B, and 4A). At the transcript level, it was shown that both *LhcB2* and *LhcB2-GFP* were up regulated by 24-h illumination in all three lines (Figure 5C).

**Figure 5.**
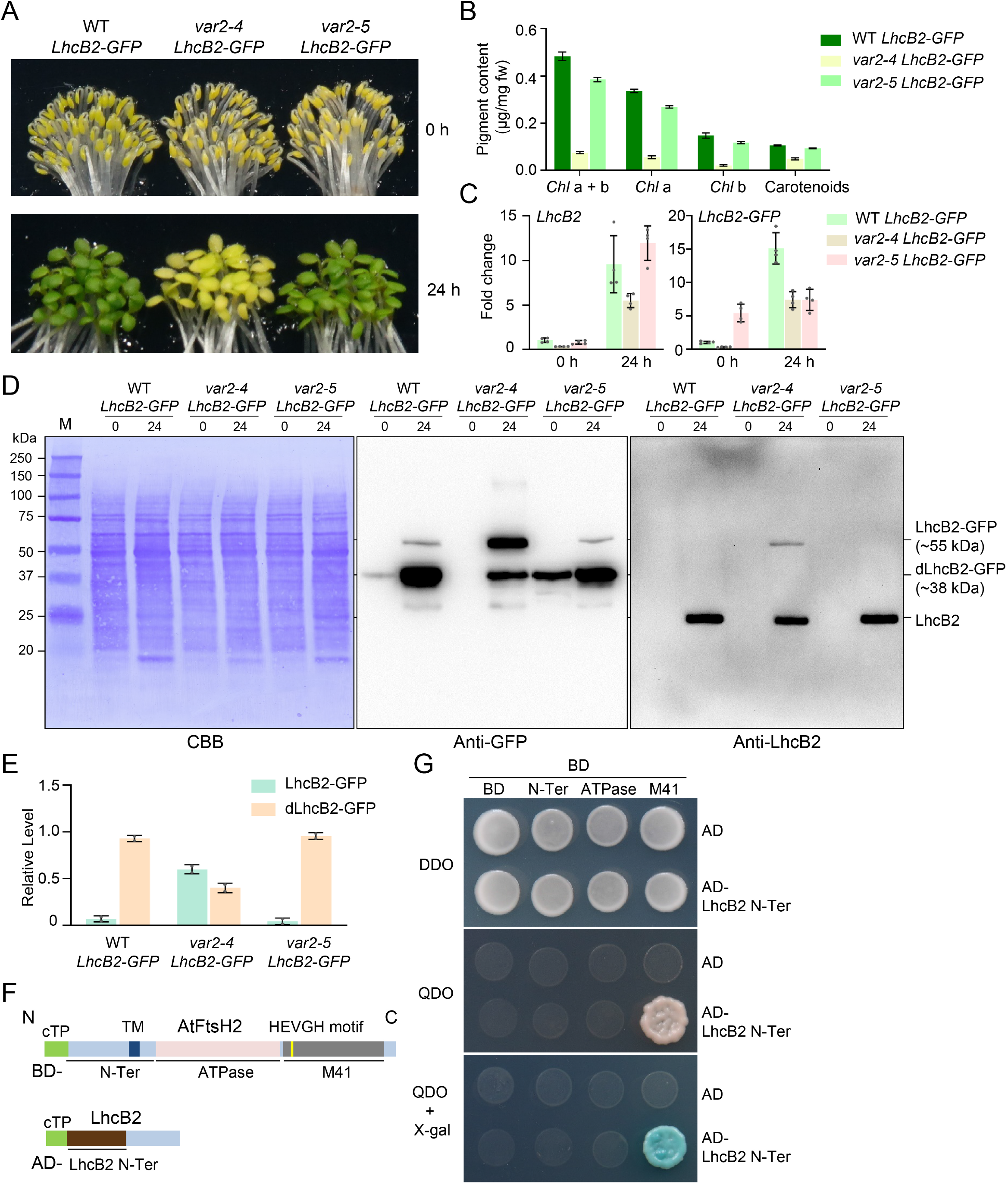
Thylakoid FtsH is required for the N-terminal degradation of LhcB2-GFP during de-etiolation. A. Phenotypes of etiolated and 24-h de-etiolated WT *LhcB2-GFP*, *var2-4 LhcB2-GFP*, and *var2-5 LhcB2-GFP*. B. The accumulation of photosynthetic pigments in 24-h de-etiolated seedlings shown in (A). C. The accumulation of *LhcB2* and *LhcB2-GFP* transcripts in etiolated and 24-h de-etiolated seedlings shown in (A). qRT-PCR analyses were performed, and fold changes were calculated using *PP2A* as the reference gene. Data are means ± s.d. of four biological replicates. D. The accumulation of LhcB2-GFP and dLhcB2-GFP in 24-h de-etiolated WT *LhcB2-GFP*, *var2-4 LhcB2-GFP*, and *var2-5 LhcB2-GFP*. Protein loadings were normalized to equal fresh tissue weight, and confirmed by a Coomassie Brilliant Blue-stained PVDF membrane (CBB). E. Quantitative analyses of the relative level of LhcB2-GFP and dLhcB2-GFP in (D). The total signal intensity of LhcB2-GFP and dLhcB2-GFP detected by the anti-GFP antibody, was defined as 1.0 in each genotype. Scatter plots with error bars are from two biological repeats. F. Schematic representation of the N-terminus of LhcB2 (LhcB2 N-Ter, used as a prey, AD-) and domains of VAR2/AtFtsH2 (used as baits, BD-) for yeast two-hybrid assays. The LhcB2 N-Ter was deduced from the relative size of dLhcB2-GFP. cTP for chloroplast transit peptide, TM for the putative transmembrane domain, ATPase for the ATPase domain, and M41 for the zinc-dependent protease domain of VAR2/AtFtsH2. G. The M41 domain of VAR2/AtFtsH2 interacts with the N-terminus of LhcB2 in yeasts. QDO plates supplemented with X-α-Gal for a higher stringency.

We further analyzed the accumulation of LhcB2-GFP and dLhcB2-GFP in 24-h de-etiolated WT *LhcB2-GFP*, *var2-4 LhcB2-GFP*, and *var2-5 LhcB2-GFP*. Less dLhcB2-GFP and more LhcB2-GFP were accumulated in 24-h de-etiolated *var2-4 LhcB2-GFP* compared with those in WT *LhcB2-GFP* and *var2-5 LhcB2-GFP* (Figure 5D). The relative accumulation ratio of dLhcB2-GFP and LhcB2-GFP were further quantified using the immunoblotting signal intensities (Figure 5E). To further confirm thylakoid FtsH is involved in the N-terminal degradation of LhcB2-GFP, we performed yeast two-hybrid assays using the N-terminus of LhcB2-GFP as a prey, and different domains of VAR2/AtFtsH2 as baits (Figure 5F). It was shown that N-terminus of LhcB2-GFP interacted with the M41 protease domain of VAR2/AtFtsH2 in yeasts (Figure 5G). These together indicated that thylakoid FtsH is the most likely protease involved in the degradation of LhcB2-GFP to dLhcB2-GFP.

### The ATPase domain determines the degradation of LhcB2-GFP to dLhcB2-GFP

The AAA family proteases use ATPase domains to unfold substrate protein for sequential degradation achieved by its protease domains. All reported amino acid substitution mutation in *var2* alleles occurs in the ATPase domain of VAR2/AtFtsH2 (Sakamoto et al., 2004; Zhang et al., 2010). Although both *var2-5* (P320L) and *var2-3* (G267D) contain missense mutations in the ATPase domain of VAR2/AtFtsH2, *var2-3* showed a more severe leaf variegation phenotype compared with *var2-5* (Chen et al., 2000). This prompted us to investigate the role of the ATPase domain in the degradation of LhcB2-GFP using *var2-5* and *var2-3* (Figure 6A). At 24-h de-etiolation, the *var2-3* mutant showed a conspicuous pale-green phenotype, resembled to the null mutant *var2-4* (Figure 6A). We further analyzed the accumulation of VAR2/AtFtsH2 in those *var2* alleles during 24-h de-etiolation (Figure 6B). Consistent with previous findings in rosette leaves (Chen et al., 2000), all three *var2* alleles had reduced level of VAR2/AtFtsH2 compared with WT at 24-h de-etiolation (Figure 6B). Although *var2-3* contained a relatively higher level of VAR2/AtFtsH2 than *var2-5*, it showed a more severe pale-green phenotype than *var2-5* during 24-h de-etiolation (Figure 6A and B). This indicated that AtFtsH2^G267D^ in *var2-3* was less active compared with AtFtsH2^P320L^ in *var2-5*. In *var2-3 LhcB2-GFP* at 24-h de-etiolation, a higher ratio of LhcB2-GFP/dLhcB2-GFP was also observed, resembled to that in *var2-4 LhcB2-GFP* (Figure 6C and D, and Supplemental Figure 5). This indicated that the ATPase domain of VAR2/AtFtsH2 determines the degradation of LhcB2-GFP to dLhcB2-GFP.

**Figure 6.**
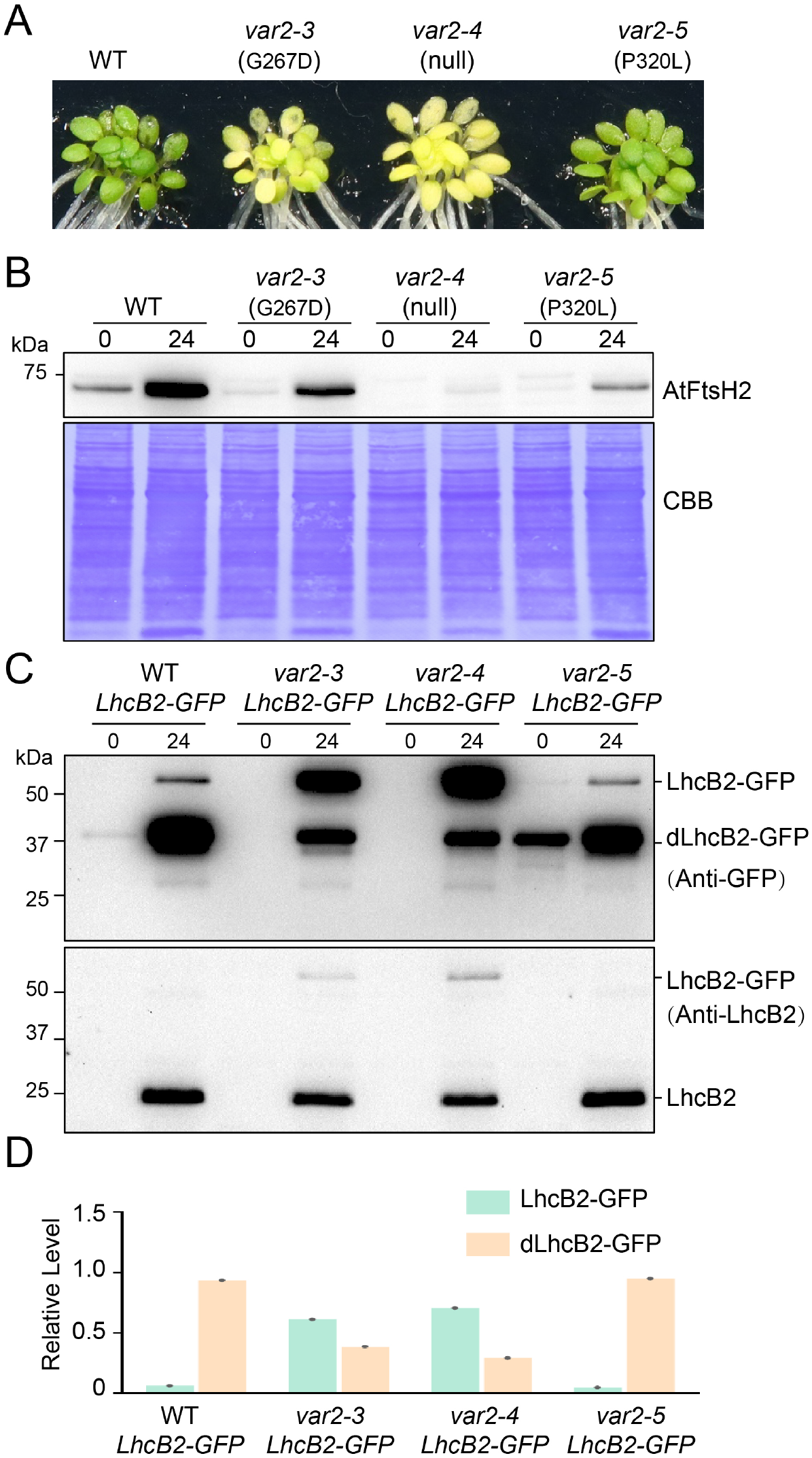
The ATPase domain is required for the degradation of LhcB2-GFP. A. The phenotype of 24-h de-etiolated WT, *var2-3, var2-4*, and *var2-5*. B. The accumulation of VAR2/AtFtsH2 in 24-h de-etiolated WT, *var2-3, var2-4*, and *var2-5*. C. The accumulation of LhcB2-GFP and dLhcB2-GFP in 24-h de-etiolated WT *LhcB2-GFP*, *var2-3 LhcB2-GFP*, *var2-4 LhcB2-GFP*, and *var2-5 LhcB2-GFP*. The uncropped original figures were shown in Supplemental Figure 5. D. Quantitative analyses of relative levels of LhcB2-GFP and dLhcB2-GFP in (C).

### The accumulated LhcB2-GFP in *var2-3* and *var2-4* also form aggregates

We next examined the accumulation of LhcB2-GFP in cotyledons of *var2-3 LhcB2-GFP*, *var2-4 LhcB2-GFP* and *var2-5 LhcB2-GFP in vivo* (Figure 7A). In WT *LhcB2-GFP*, the GFP fluorescence was dispersed in chloroplasts and overlap with the chlorophyll fluorescence, indicating the chloroplast localization of LhcB2-GFP and dLhcB2-GFP (Figure 7A). Consistent with those observed in *pga4 LhcB2s*, GFP fluorescence was detected as discrete foci surrounding chlorophyll fluorescence in 24-h de-etiolated *var2-3 LhcB2-GFP* and *var2-4 LhcB2-GFP*, (Figure 7A and 4E). However, little foci were observed in the weaker allele *var2-5 LhcB2-GFP* (Figure 7A).

**Figure 7.**
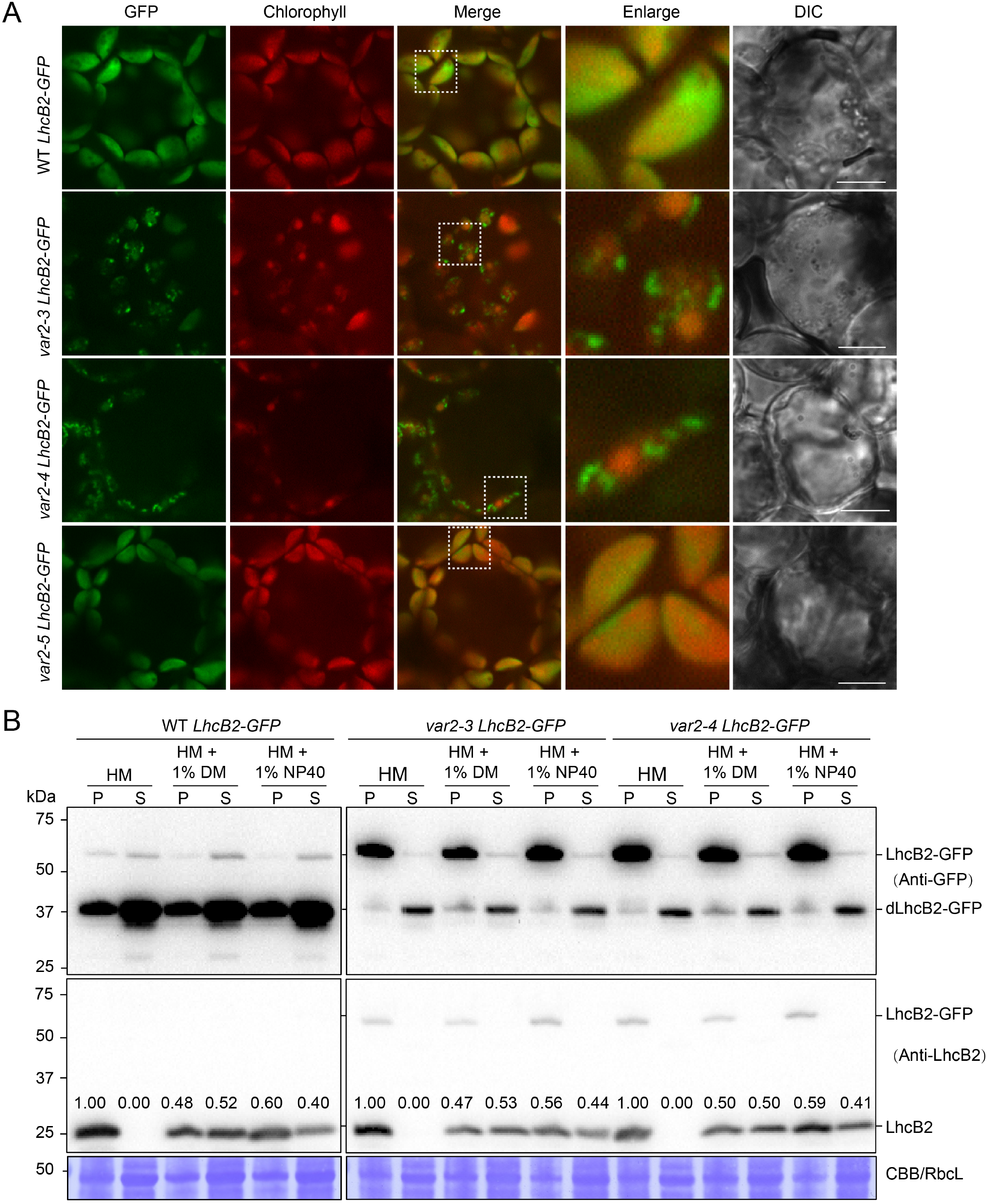
The accumulated LhcB2-GFP in *var2-3* and *var2-4* form aggregates. A. The detection of GFP and chlorophyll fluorescence in 24-h de-etiolated WT *LhcB2-GFP*, *var2-3 LhcB2-GFP*, *var2-4 LhcB2-GFP*, and *var2-5 LhcB2-GFP*. DIC for differential interference contrast images. White boxes indicated enlarged areas. Bars, 10 μm. B. The solubility of LhcB2-GFP in 24-h de-etiolated WT *LhcB2-GFP*, *var2-3 LhcB2-GFP*, and *var2-4 LhcB2-GFP*. Two mild nonionic detergents n-dodecyl-β-D-maltoside (1% β-DM) and Nonidet P-40 (1% NP40) were used to solubilize membrane proteins. The anti-GFP antibody and the anti-LhcB2 antibody were used, respectively. The accumulation of LhcB2 in the P and S fractions was relatively quantified with signal intensities.The stroma protein RbcL was shown by CBB-stained PVDF membrane.

To reveal the nature of these GFP aggregates, we tried chloroplast fractionation using 24-h de-etiolated seedlings of WT *LhcB2-GFP, var2-3 LhcB2-GFP*, and *var2-4 LhcB2-GFP* (Figure 7B). In WT *LhcB2-GFP*, LhcB2 was present in the P fraction, or partially solubilized in the S fraction by using either 1% β-DM or 1% NP-40 (Figure 7B). Meanwhile, a large amount of dLhcB2-GFP were present in the S fraction in the HM buffer or HM buffer containing detergents, indicating that they were mainly in the soluble stroma fraction (Figure 7B and 3B). Consistently, partial LhcB2-GFP was solubilized when nonionic detergents were used in WT *LhcB2-GFP*, indicating LhcB2-GFP was associated with thylakoid membranes (Figure 7B and 3B). In contrast, most of the accumulated LhcB2-GFP in *var2-3 LhcB2-GFP* and *var2-4 LhcB2-GFP* were resistant to nonionic detergents (Figure 7B). There were still some dLhcB2-GFP and endogenous LhcB2 were not solubilized by nonionic detergents (Figure 7B and 3B). These together indicated that LhcB2-GFP are prone to form protein aggregates in both *var2 LhcB2-GFPs* and *pga4 LhcB2-GFPs*. Interestingly, the ratio of detergent-dissolved LhcB2 to detergent-resistant LhcB2 was not affected in *var2-3 LhcB2-GFP* or *var2-4 LhcB2-GFP* compared with those in the WT *LhcB2-GFP* (Figure 7B).

### Other chloroplast proteases are involved in accumulation of LhcB2-GFP

Chloroplasts contain numerous proteases for chloroplast PQC (Nishimura et al., 2017; Sun et al., 2021). We checked if other chloroplast proteases are involved in the accumulation of LhcB2-GFP and degradation of LhcB2-GFP to dLhcB2-GFP, including the ATP-independent metalloprotease EGY1 (Chen et al., 2005), a putative thylakoid-localized M41 metalloprotease VIR3 (Qi et al., 2016), the chloroplast envelope-localized AtFtsH12 (Kikuchi et al., 2018), and the stroma-localized Clp serine protease (Bouchnak and van Wijk, 2022). Accordingly, we crossed WT *LhcB2-GFP* with related mutants *egy1-3, vir3-1, pga1-1* (a hypomorphic allele of *AtFtsH12*) (Li et al., 2022), and *clpr1-2* (Koussevitzky et al., 2007). During 24-h de-etiolation, the accumulation of LhcB2-GFP and dLhcB2-GFP varied in those mutants (Supplemental Figure 6). However, ratios of LhcB2-GFP/dLhcB2-GFP were almost unaffected in those mutants compared with that in WT *LhcB2-GFP* (Supplemental Figure 6). Interestingly, the accumulation of both LhcB2-GFP and dLhcB2-GFP was severely reduced in *pga1-1* and *clpr1-2*, suggesting their roles in chloroplast protein homeostasis (Li et al., 2022).

### *PGA4* is a new suppressor locus of *var2* leaf variegation

As both FtsH and cpSRP54 are involved in the degradation of LhcB2-GFP, we analyzed the genetic interaction between *pga4-1* and *var2* mutants in this process. Moreover, the *pga4-2* mutant was obtained from an activation tagging mutagenesis pool of *var2-5*, indicated that *PGA4* is likely a new suppressor locus for the leaf variegation phenotype of *var2*. The single mutant *pga4-1* showed pale-green rosette leaves, while *var2* mutants showed variegated rosette leaves (Figure 8A-C). The double mutant *var2-5 pga4-1* was obtained, in which the leaf variegation of *var2-5* was masked by *pga4-1* (Figure 8A). In addition, *pga4-1* could also suppress *var2-4* (Figure 8B). The binary vector *ProcpSRP54:cpSRP54-GFP* was used to recover the double mutant *var2-4 pga4-1* to *var2-4* (Figure 8C). These confirmed that the *pga4-1* mutant is a new genetic suppressor of *var2*.

**Figure 8.**
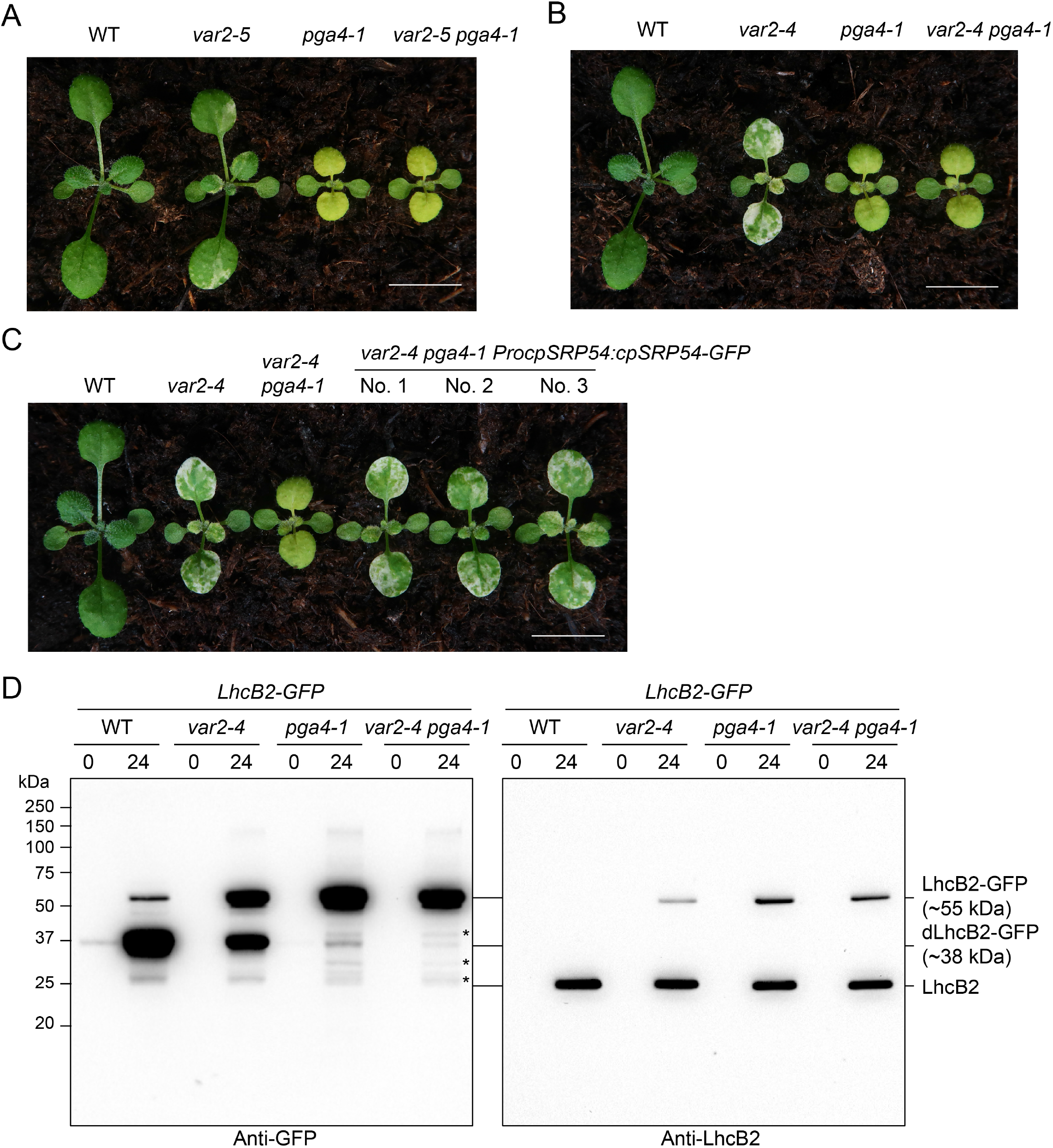
The *pga4-1* mutant is a new *var2* suppressor. A. Representative WT, *var2-5, pga4-1* and *var2-5 pga4-1*. B. Representative WT, *var2-4, pga4-1* and *var2-4 pga4-1*. C. Complementation of the double mutant *var2-4 pga4-1* to *var2-4* with the binary construct *ProcpSRP54:cpSRP54-GFP*. Representative two-week-old seedlings were shown. Bars, 1.0 cm. D. The accumulation of LhcB2-GFP and dLhcB2-GFP in 24-h de-etiolated WT *LhcB2-GFP*, *var2-4 LhcB2-GFP*, *pga4-1 LhcB2-GFP* and *var2-4 pga4-1 LhcB2-GFP*. Asterisks indicated other possible degraded LhcB2-GFP products in *pga4-1 LhcB2-GFP* and *var2-4 pga4-1 LhcB2-GFP*.

We further checked the accumulation of LhcB2-GFP and dLhcB2-GFP in the double mutant *var2-4 pga4-1 LhcB2-GFP* during de-etiolation. The accumulation of dLhcB2-GFP was even more reduced in *var2-4 pga4-1 LhcB2-GFP* when compared with those in *pga4-1 LhcB2-GFP* (Figure 8D). This suggested that *PGA4* is epistatic to *VAR2/AtFtsH2* in the degradation of LhcB2-GFP and the formation of leaf variegation. Interestingly, we also observed that some other faint degraded products of LhcB2-GFP, which became relatively obvious in *pga4-1 LhcB2-GFP* and *var2-4 pga4-1 LhcB2-GFP* (Figure 8D). The accumulation of those degraded products was comparable between *pga4-1 LhcB2-GFP* and *var2-4 pga4-1 LhcB2-GFP*, indicating the existence of other proteases to clean LhcB2-GFP aggregates.

Taken together, we presented a model that cpSRP54 and thylakoid FtsH could determine the N-terminal degradation of LhcB2-GFP sequentially on thylakoid membrane (Figure 9). In WT *LhcB2-GFP*, the heterogeneous LhcB2-GFP will be targeted to thylakoid membranes predominantly by cpSRP54-dependent protein translocation pathway (1). The chimeric LhcB2-GFP is not assembled into functional complexes and subjected to N-terminal degradation mediated by thylakoid FtsH. The FtsH degrades LhcB2-GFP, initiated at the N-terminus and ceased before approaching the stable structure of GFP protein, and releases the dLhcB2 into stroma (2). The ATPase domain of FtsH provides essential driving forces to unfold LhcB2-GFP for further degradation performed by the protease domain. Alternatively, the lower abundance of dLhcB2-GFP in the *SRP54* null mutant *pga4-1 LhcB2-GFP* indicated that a minor fraction of LhcB2-GFP is transported to thylakoid membranes by cpSRP54-independent pathways. The stroma localized LhcB2-GFP and dLhcB2 may forms protein aggregates, resistant to mild nonionic detergents (3 and 4). In *pga4 LhcB2-GFPs* or *var2 LhcB2-GFPs*, the mutation of either cpSRP54-dependent translocation, or the activity of thylakoid FtsH could prevent the degradation of LhcB2-GFP to dLhcB2, which in turn form protein aggregates. Also, there are possibilities that some stromal chaperones or proteases would be involved in the fully degradation of LhcB2-GFP and dLhcB2-GFP to free amino acids to maintain chloroplast protein homeostasis (5).

## Discussion

### Using the unassembled chimeric LhcB2-GFP as a heterogeneous marker to understand translocation of thylakoid membrane proteins

The light-harvesting chlorophyll a/b binding (LHC) proteins, encoded by the nuclear genome, are the most abundant thylakoid membrane proteins (Schünemann, 2004). The expression of *LHCPs* was used as marker genes in understanding of the anterograde and retrograde regulation of chloroplast development (Susek et al., 1993; Jarvis and Lopez-Juez, 2013). To understand the chloroplast protein homeostasis, we systematically isolation *pale green Arabidopsis* (*pga*) mutants (Li et al., 2022). In this work, we reported that *PGA4* encodes cpSRP54 (Figure 1). To further dissect the function of cpSRP54 in chloroplast protein homeostasis, we generated a stable transgenic marker line to express a chimeric *LhcB2-GFP* resembling to the endogenous *LhcB2* (Figure 2). Our biochemical assays showed that the chimeric LhcB2-GFP is targeted into thylakoid membranes during de-etiolation (Figure 3). More interestingly, the membrane-associated LhcB2-GFP is not able to be incorporated into LHCII complexes for unknown reasons, and further degraded to a predominant form dLhcB2-GFP in WT *LhcB2-GFP* (Figure 2 and 3). Although the C terminal fusion with GFP may disturb the proper assemble of LhcB2-GFP into functional LHCII complexes, this chimeric LhcB2-GFP and the degraded form dLhcB2-GFP provided a heterogeneous unassembled protein marker for protein translocation to thylakoids. Using *pga4 LhcB2-GFPs*, we demonstrated that both cpSRP54-dependent and cpSRP54-independent translocation pathways exist in cotyledons, and the former might be functional predominantly for LhcB2-GFP.

Surprisingly, the degraded form dLhcB2-GFP is sharply reduced in *pga4-1 LhcB2-GFP* compared with those in WT *LhcB2-GFP* (Figure 4). Moreover, the accumulated LhcB2-GFP formed insoluble protein aggregates and discrete GFP fluorescent foci in *pga4* mutants. Without chlorophyll pigments, some of those accumulated LhcB2 could be aggregated apoproteins, of which proportion being insoluble to mild detergents in *pga4-1 LhcB2-GFP* is relatively higher than those in WT *LhcB2-GFP*. These provided *in vivo* evidences that cpSRP54 could prevent aggregations of LHC apoproteins (Schünemann et al., 1998; Yuan et al., 2002; Goforth et al., 2004). The translocation of unassembled LhcB2-GFP to thylakoid is a prerequisite for its thylakoid-associated degradation to dLhcB2-GFP, and this scenario may explain the discrepancy in accumulations of photosynthetic pigments and endogenous LhcB2 during de-etiolation in *pga4-1 LhcB2-GFP*. However, the degradation of insoluble endogenous LhcB2 in *pga4-1 LhcB2-GFP* requires further experimental supports, as we could not observe any LhcB2 C terminal fragments with the LhcB2 antibody. Although we did not have direct evidences to support the degradation of LhcB2-GFP is consistent with the endogenous LhcB2, the discrete LhcB2-GFP fluorescent foci in *pga4-1 LhcB2-GFP* could be used as a heterogeneous marker to dissect the regulatory network in chloroplast PQC that interconnecting with cpSRP54.

### Using the degradation of LhcB2-GFP to dLhcB2-GFP to dissect chloroplast PQC during de-etiolation

FtsH is a prevailing membrane-localized AAA family protease that can degrade misfolded or unassembled polypeptides for membrane protein quality control in prokaryotes and organelles of eukaryotes (Janska et al., 2013). In photosynthetic species, it is generally accepted that the photo-damaged core protein D1 of photosystem II complexes is a bona fide substrate of thylakoid FtsH proteases (Nixon et al., 2010; Kato et al., 2012). However, our understanding on FtsH-mediated thylakoid membrane PQC in higher plants during de-etiolation is still limited. The *var2* mutants, with green cotyledons when grown under normal conditions, showed conspicuous delayed de-etiolation phenotypes, indicating indispensible roles of FtsH in thylakoid membrane PQC during de-etiolation. Our genetic and biochemical evidences strongly supported that thylakoid FtsH is involved in the degradation of LhcB2-GFP to dLhcB2-GFP during de-etiolation. The use of different *var2* alleles further indicated that the ATPase activity of thylakoid FtsH determines the degradation of LhcB2-GFP.

We showed that thylakoid FtsH degrades the unassembled LhcB2-GFP to dLhcB2-GFP, providing a trackable marker protein to dissect FtsH-mediated thylakoid membrane PQC during de-etiolation. The degradation of this heterogeneous substrate LhcB2-GFP supported that cpSRP54 and thylakoid FtsH complex work sequentially for thylakoid membrane protein quality control (Figure 9). Similarly to those in *pga4-1 LhcB2-GFP*, endogenous LhcB2 was highly accumulated in *var2-4 LhcB2-GFP* and *var2-4 pga4-1 LhcB2-GFP*. Thus it is reasonable to assume that the rapid turnover of misfolded or unassembled subunits is essential to avoid the formation of toxic aggregates during de-etiolation, because thylakoid membrane-localized photosynthetic proteins are mostly hydrophobic. Moreover, we observed that the degradation of LhcB2-GFP to other types of fragments in *pga4-1 LhcB2-GFP* and *var2-4 pga4-1 LhcB2-GFP*, supporting that some stroma localized chaperones or proteases were also involved in degrading LhcB2-GFP or dLhcB2-GFP aggregates. It was observed that the accumulation of Clp, CPN60 and cpHSP70 were increased in *cpspr54/ffc* or *var2* mutants (Amin et al., 1999; Rutschow et al., 2008; Dogra et al., 2019). These may participate in the degradation of misfolded/unassembled membrane proteins that accumulated in the stroma, possibly compensating the loss of cpSRP54 and FtsH.

**Figure 9.**
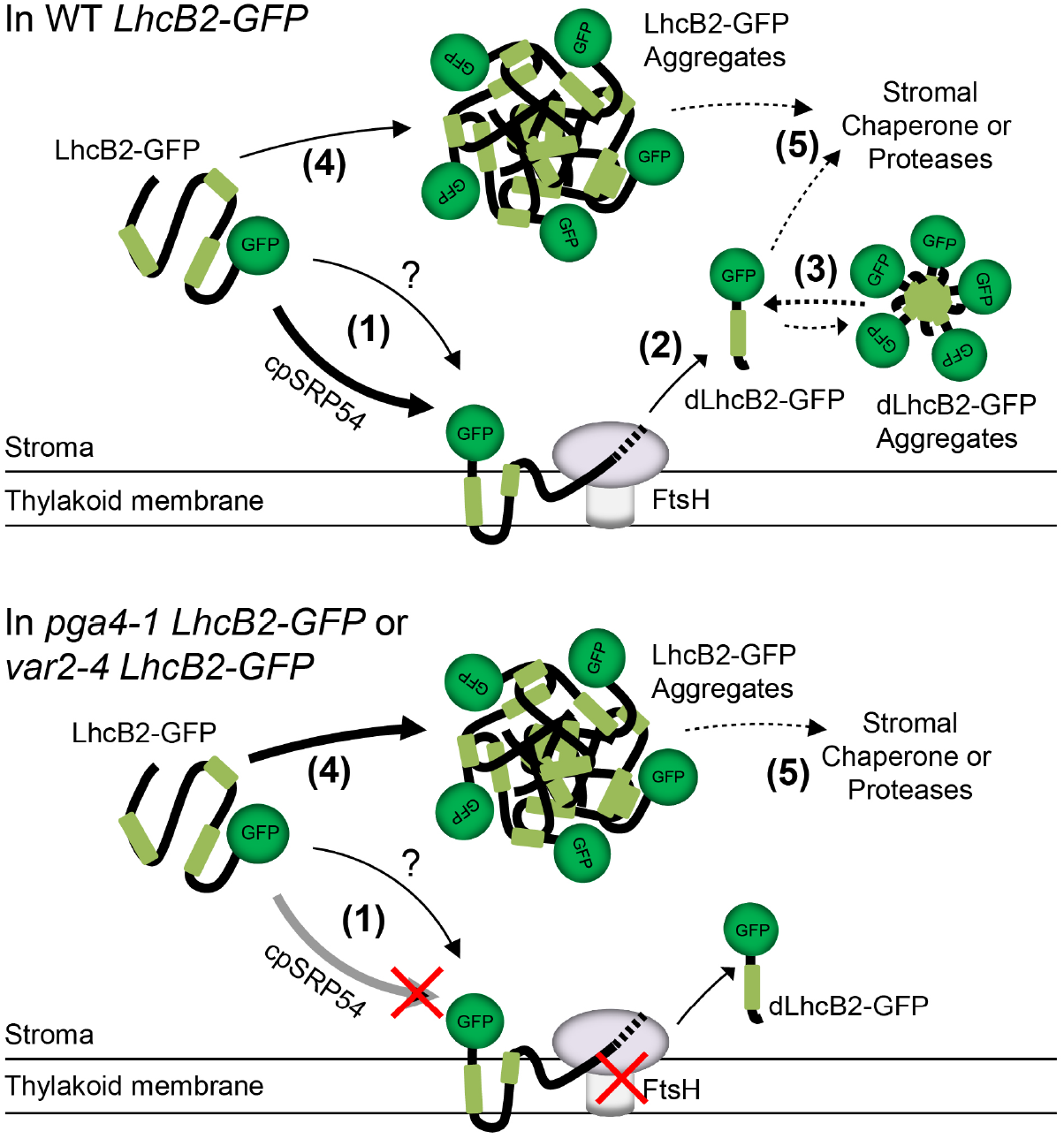
A working model for cpSRP54 and thylakoid FtsH modulate N-terminal degradation of LhcB2-GFP. In WT *LhcB2-GFP*, LhcB2-GFP is translocated to the thylakoid membrane via the SRP54-dependent pathway (1), and degraded by the thylakoid FtsH. The question mark indicates SRP54-independent translocation pathways. When encountering the stable GFP structure, FtsH stops degradation, and releases dLhcB2-GFP either to stroma (2). The released dLhcB2-GFP may forms dLhcB2-GFP aggregates (3). In *pga4-1 LhcB2-GFP* or *var2-4 LhcB2-GFP*, disruption of either the cpSRP54-dependent translocation or the thylakoid FtsH activity, could prevent the degradation of LhcB2-GFP to dLhcB2 on thylakoid membrane, which in turn form LhcB2-GFP aggregates in the stroma (4). Other chaperones or proteases may involve in the degradation of LhcB2-GFP or dLhcB2-GFP aggregates in the stroma (5).

### The partial degraded LhcB2-GFP indicates the N-terminal processing activity of thylakoid FtsH

The most characterized degradation process mediated by thylakoid FtsH is the D1 degradation during PSII repair cycle under high light conditions. FtsH is required for the processive degradation of the full D1 or the N terminal fragments of D1 generated by Deg proteases (Komenda et al., 2007; Kato et al., 2012; Kato et al., 2018). *var2* mutants contain high level of full D1 under high light stress (Bailey et al., 2002; Kato et al., 2012; Qi et al., 2020; Wang et al., 2022). In cyanobacteria, the N terminal D1 sequence and FtsH proteases determine the stability of D1 (Komenda et al., 2007). Similar to D1 degradation, the degradation of LhcB2-GFP to dLhcB2-GFP is initiated at its N terminus, but ceased before approaching the stable structure of GFP protein. Thus, the accumulation of LhcB2-GFP and dLhcB2-GFP provided easy-trackable substrate and product to analyze the N-terminus degradation thylakoid FtsH under nonphotoinhibitory light conditions for future studies.

Instead of being fully degraded, the accumulation of dLhcB2-GFP fragments raises a fundamental question that if thylakoid FtsH proteases have protein processing activity at physiological conditions (Okuno and Ogura, 2013). Other FtsH homologs from *E.Coli* and yeast mitochondria, can not only degrade one entire protein processively, but also perform protein modifications by protein N or C terminal processing *in vitro* or *in vivo* (Okuno et al., 2006; Walker et al., 2007; Bonn et al., 2011; Chauleau et al., 2011). The remained dLhcB2-GFP demonstrated the potential N-terminal processing activity of thylakoid FtsH, which is consistent with their prokaryotic homologs. If this N-terminal processing activity is conserved in the thylakoid FtsH, one plausible scenario is that thylakoid FtsH is required for processing, instead of full degradation of certain proteins that are essential for thylakoid membrane biogenesis and chloroplast development. Although the FtsH is one of the best characterized AAA family protease, we still know little about its potential substrates during de-etiolation, and how it determines chloroplast development during de-etiolation. Future work on the identification of thylakoid FtsH substrates, will address this point.

### Reduced thylakoid membrane protein loading may alleviate the *var2* leaf variegation phenotype

One intriguing feature of *var2* mutants is the variegated rosette leaves with normal-appearing green sectors, and albino sectors with abnormal chloroplasts. Systematic identification of genetic modifiers of the variegation phenotype of *var2*, unraveled numerous loci that interact with *VAR2*/*AtFtsH2* genetically. The identification of SVR8/RPL24, SVR9/IF3, and SVR11/EF-Tu, indicated that weakened chloroplast translation could suppress the leaf variegation of *var2* (Liu et al., 2013; Zheng et al., 2016; Liu et al., 2019). Nonetheless, the molecular mechanism is still not well understood. Here, we found that *pga4* mutants are new *var2* suppressors (Figure 8). Besides the translocation of LHC proteins to thylakoid membranes, it was reported recently that cpSRP54 interacts with chloroplast ribosomes to navigate the insertion of *de novo* synthesized nascent proteins, such as the PSII reaction center protein D1 (Ziehe et al., 2018; Hristou et al., 2019). Given the fact that a group of SVRs are involved in chloroplast translation, it was deduced that the defects in cpSRP54-mediated translocation of chloroplast translated nascent proteins caused suppression of the *var2* leaf variegation. In addition, D1 is vulnerable to light and photo-damaged, and the thylakoid FtsH is involved in the degradation of photo-damaged D1 (Nixon et al., 2010). The identification of *pga4-1* provided us a scenario that cpSRP54 and thylakoid FtsH complex may work coordinately to the membrane protein loading and degradation balance, especially two most abundant pigment-binding photosynthetic proteins LHCP and D1, and this membrane protein homeostasis may further influence the leaf variegation phenotype and chloroplast development defects in *var2* mutants.

## Materials and Methods

### Plant materials and growth conditions

All *Arabidopsis thaliana* strains used in this study are the Columbia ecotype (Col), except for the Landsberg *erecta* ecotype (L*er*) to generate the mapping population. Arabidopsis mutants *var2-3, var2-4, var2-5, pga1-1, vir3-1, evr3-1, clpr1-2* are described in results. The *pga4-1* mutant was isolated from an ethyl methanesulfonate mutagenesis pool, and backcrossed with Col for 7 times. The *pga4-2* mutant was recovered from an activation tagging mutagenesis pool in *var2-5* background, and backcrossed with Col for 3 times (Yu et al., 2008). Seeds were surface sterilized and stratified at 4 °C for 2 d before planting. For de-etiolation assay, sterilized seeds were grown on 1/2 Murashige and Skoog (1/2 MS) media containing 1% Bacto agar in the dark for 3 d at 22 °C, and transferred to continuous light (~100 μmol m^-2^ s^-1^ photons) in a growth chamber for indicated time periods. For chemical treatments, seedlings were grown on 1/2 MS media containing 0.5 mM lincomycin (Lin) or 5 μM norflurazon (NF), respectively. Plants for others purpose were grown on commercial soil mix (Pindstrup) under continuous light (~100 μmol m^-2^ s^-1^ photons) at 22 °C.

### Map-based cloning of *PGA4*

Map-based cloning was performed to identify the *PGA4* locus. The *pga4-1* mutant was crossed with the L*er* ecotype to generate the *F2* mapping population. Bulked segregant analysis located *PGA4* to a region near to the marker *F8F6#1* on chromosome 5. Additional molecular markers based on Indel or SNP polymorphisms were designed for fine mapping.

### Construction of vectors and generation of transgenic lines

Genomic sequences of *AtLhcB2.2* with endogenous promoter regions (2089 bp), and genomic sequences of *cpSRP54* with endogenous promoter regions (1353 bp), were cloned into the binary vector *pCambia1300-GFP*, to generate *ProLhcB2:gLhcB2-GFP* and *ProcpSRP54:cpSRP54-GFP*. Transgenic lines were generated by the *Agrobacterium tumefaciens*-mediated floral dip method (Clough and Bent, 1998). Transgenic lines were screened on 1/2 MS media containing 25 mg L^-1^ hygromycin. The *var2* or *pga4 LhcB2-GFP* lines were obtained by genetic crosses with WT *LhcB2-GFP*, and *F3* generations carrying the hygromycin resistant marker without segregation were used for experiments. Primers for vectors and genotyping are listed in Supplemental Table 1.

### RNA extraction and quantitative RT-PCR (qRT-PCR)

Total RNAs were extracted from de-etiolated seedlings using the TRIzol reagent (15596026, Thermo Fisher Scientific). cDNAs were synthesized from 1.0 μg total RNA with the UEIris RT mix with DNase (All-in-one) (R2020, US Everbright). qPCRs were performed using 2 × ChamQ SYBR qPCR Master Mix (Q311-02, Vazyme). Primers for qRT-PCRs are listed in Supplemental Table 1.

### Protein extraction and immunoblotting analyses

The separation of soluble and membrane fractions were performed as described (Hou et al., 2021), with some modifications. De-etiolated seedlings were weighed and ground in HM buffer (10 mM Hepes-KOH pH7.6, 5 mM MgCl_2_), with or without 2 M NaCl, 1% n-dodecyl-β-D-maltoside or 1% Nonidet P-40 if required. Supernatant (S) and pellet (P) were separated by centrifugation at 20,000 g for 15 min at 4 °C. Pellet was further resuspended with the same volume of HM buffer. Both S and P fractions were diluted with the same volume of 2 × SDS sample buffer (0.125M Tris-HCl pH6.8, 4% SDS and 20% glycerol, 1% β-mercaptoethanol) for further SDS-PAGE and immunoblotting. To extract total proteins, fresh plant seedlings were weighed and ground in liquid nitrogen, and incubated with 2 × SDS sample buffer at 65 °C for 1 h. After centrifugation at 18,000 g for 10 min, the supernatant were collected and loaded on 12% SDS-PAGE containing 8M urea for electrophoresis. Protein loading was normalized to equal fresh tissue weight. After transferred to PVDF membranes (0.22 μm, Millipore), immunoblotting analyses were performed with standard protocols. The anti-GFP antibody (TaKaRa, 632381), the anti-LhcB2 antibody (Agrisera, AS01003), the anti-VAR2/AtFtsH2 antibody (Qi et al., 2016), and the anti-RbcL antibody (Agrisera, AS03037) were used. The quantification of signal intensity was carried out by the Image Lab software (Bio-Rad).

### Blue Native PAGE

Preparation of BN-PAGE gel was performed as described (Järvi et al., 2011). De-etiolated WT *LhcB2* seedlings were weighed and homogenized in liquid nitrogen. HM buffer was used to suspend the homogenate, and Supernatant (S) and pellet (P) were separated by centrifugation at 20,000 g for 15 min at 4 °C. Thylakoid membranes from the pellet fraction were solubilized with 1% β-DM (w/v) in 25BTH20G buffer (25 mM Bis-Tris-HCl pH7.0 and 20% glycerol). The β-DM solubilized fraction was collected by centrifugation at 20,000 g for 10 min at 4 °C, and resolved on 3-10% 1-D BN-PAGE gel. Excised gel lanes from the 1-D BN-PAGE gel were incubated in 2 × SDS sample buffer with gentle shaking at room temperature for 30 min for denaturation. The 1-D gel was further applied to 10% SDS-PAGE gel for 2-D electrophoresis.

### Recombinant protein expression in *Escherichia coli*

The coding regions for LhcB2, ΔN-LhcB2, LhcB2-ΔC were cloned into the pGEX4T-1 vector. Expressions of GST-LhcB2, GST-ΔN-LhcB2, GST-LhcB2-ΔC in *Escherichia coli* BL21 (DE3) were induced with 0.1 mM IPTG at 37 °C for 4 h. Primers for the vector construction are listed in Supplemental Table 1.

### Quantification of chlorophyll content

To measure the chlorophyll content, de-etiolated seedlings were harvested, weighted and ground in liquid nitrogen. Chlorophyll was extracted by 95% ethanol (v/v) with gentle rotation for 24 h at 4 °C. Tissue debris was removed by centrifugation at 19,000 g for 20 min at 4 °C. The content of chlorophyll and carotenoids was calculated as described (Wang et al., 2018).

### Yeast two-hybrid assays

The yeast strain Y2HGold and the Matchmaker Gold Yeast Two-Hybrid System (Clontech) were used for the yeast two-hybrid assay. The coding region for the N-terminus of LhcB2 without chloroplast transit peptide (amino acid residues 42 to 158) was cloned into pGADT7 (the AD vector for prey). The coding regions for the N-terminal domain (amino acids residues 47-220), the ATPase domain (amino acids residues 221-466), and the M41 protease domain (amino acids residues 473-671) of VAR2/AtFtsH2 were cloned into pGBKT7 (the BD vector for bait). The AD- and BD- vectors were co-transformed into Y2HGold cells, and selected on the double dropout SD medium (DDO, lacking leucine and tryptophan). Protein interactions were verified by grown colonies on the quadruple dropout SD medium (QDO, lacking adenine, histidine, leucine, and tryptophan), and QDO medium supplemented with 100 μg mL^-1^ X-α-gal for high stringency. Primers are listed in Supplemental Table 1.

### Confocal microscopy

GFP fluorescence and chlorophyll auto-fluorescence images were collected using a spinning disk confocal microscope (Revolution-XD, Andor). GFP fluorescence was excited by the 488 nm laser, and chlorophyll auto-fluorescence by 637 nm. The *LhcB2-GFP* lines were examined after 24-h de-etiolation, and cotyledons were used directly for confocal microscopy imaging. Images were processed by the Fiji ImageJ software.

### Accession Numbers

Sequence data used in this study can be found in The Arabidopsis Information Resource (TAIR) with the following accession numbers: *LhcB2.2, AT2G05070; VAR2/AtFtsH2, AT2G30950; cpSRP54, AT5G03940; AtPP2A, AT1G69960*.

## Acknowledgments

We thank Dr. Ningjuan Fan from the Teaching and Research Core Facility at the College of Life Sciences, Northwest A&F University, for technical assistance in using confocal microscope.

## Author contributions

Y. Q. and Y. L. conceived and designed research plans; Y. Q. and F. Y. supervised the experiments; Y. L., B. L., X. W., J. W., P. W., and J. Z. performed experiments; Y. Q. and F. Y. analyzed the data; Y. L. and Y. Q. wrote the manuscript with contributions from all the authors. Y. L. and B. L. contributed equally.

## Supporting Information

This article contains supporting information.

**Supplemental Figure 1.** Isolation of two *pga4* mutants.

**Supplemental Figure 2.** Accumulation of cpSRP54-GFP in 24-h de-etiolated *pga4-1 ProcpSRP54:cpSRP54-GFP*.

**Supplemental Figure 3.** Supporting information for Figure 2.

**Supplemental Figure 4.** Supporting information for Figure 3.

**Supplemental Figure 5.** Supporting information for Figure 6.

**Supplemental Figure 6.** Accumulation of LhcB2-GFP in other chloroplast protease mutants.

**Supplemental Table 1.** Primers used in this study.

## Data availability

The data that support the findings of this study are available from the corresponding author upon reasonable request.

## References

Akopian D, Shen K, Zhang X, Shan SO (2013) Signal Recognition Particle: An Essential Protein-Targeting Machine. Annual Review of Biochemistry, Vol 82 82: 693–721

Amin P, Sy DAC, Pilgrim ML, Parry DH, Nussaume L, Hoffman NE (1999) Arabidopsis mutants lacking the 43-and 54-kilodalton subunits of the chloroplast signal recognition particle have distinct phenotypes. Plant Physiology 121: 61–70

Arlt H, Tauer R, Feldmann H, Neupert W, Langer T (1996) The YTA10-12 complex, an AAA protease with chaperone-like activity in the inner membrane of mitochondria. Cell 85: 875–885

Aro EM, Virgin I, Andersson B (1993) Photoinhibition of Photosystem II. Inactivation, protein damage and turnover. Biochim Biophys Acta 1143: 113–134

Bailey S, Thompson E, Nixon PJ, Horton P, Mullineaux CW, Robinson C, Mann NH (2002) A critical role for the Var2 FtsH homologue of Arabidopsis thaliana in the photosystem II repair cycle in vivo. J Biol Chem 277: 2006–2011

Bieniossek C, Niederhauser B, Baumann UM (2009) The crystal structure of apo-FtsH reveals domain movements necessary for substrate unfolding and translocation. Proceedings of the National Academy of Sciences of the United States of America 106: 21579–21584

Bonn F, Tatsuta T, Petrungaro C, Riemer J, Langer T (2011) Presequence-dependent folding ensures MrpL32 processing by the m-AAA protease in mitochondria. Embo Journal 30: 2545–2556

Bouchnak I, van Wijk KJ (2022) Structure, function, and substrates of Clp AAA+ protease systems in cyanobacteria, plastids, and apicoplasts: A comparative analysis (vol 296, 100338, 2021). Journal of Biological Chemistry 298

Bujaldon S, Kodama N, Rappaport F, Subramanyam R, de Vitry C, Takahashi Y, Wollman FA (2017) Functional Accumulation of Antenna Proteins in Chlorophyll b-Less Mutants of Chlamydomonas reinhardtii. Mol Plant 10: 115–130

Chauleau M, Mora L, Serba J, de Zamaroczy M (2011) FtsH-dependent Processing of RNase Colicins D and E3 Means That Only the Cytotoxic Domains Are Imported into the Cytoplasm. Journal of Biological Chemistry 286: 29397–29407

Chen G, Bi YR, Li N (2005) EGY1 encodes a membrane-associated and ATP-independent metalloprotease that is required for chloroplast development. Plant J 41: 364–375

Chen M, Choi YD, Voytas DF, Rodermel S (2000) Mutations in the Arabidopsis VAR2 locus cause leaf variegation due to the loss of a chloroplast FtsH protease. Plant Journal 22: 303–313

Clough SJ, Bent AF (1998) Floral dip: a simplified method for Agrobacterium-mediated transformation of Arabidopsis thaliana. Plant J 16: 735–743

Dogra V, Duan J, Lee KP, Kim C (2019) Impaired PSII proteostasis triggers a UPR-like response in the var2 mutant of Arabidopsis. J Exp Bot 70: 3075–3088

Glynn SE (2017) Multifunctional Mitochondrial AAA Proteases. Frontiers in Molecular Biosciences 4

Goforth RL, Peterson EC, Yuan J, Moore MJ, Kight AD, Lohse MB, Sakon J, Henry RL (2004) Regulation of the GTPase cycle in post-translational signal recognition particle-based protein targeting involves cpSRP43. J Biol Chem 279: 43077–43084

Herman C, Thevenet D, Dari R, Bouloc P (1995) Degradation of Sigma(32), the Heat-Shock Regulator in Escherichia-Coli Is Governed by Hflb. Proceedings of the National Academy of Sciences of the United States of America 92: 3516–3520

Holdermann I, Meyer NH, Round A, Wild K, Sattler M, Sinning I (2012) Chromodomains read the arginine code of post-translational targeting. Nat Struct Mol Biol 19: 260–263

Hou Z, Pang X, Hedtke B, Grimm B (2021) In vivo functional analysis of the structural domains of FLUORESCENT (FLU). Plant J 107: 360–376

Hristou A, Gerlach I, Stolle DS, Neumann J, Bischoff A, Dunschede B, Nowaczyk MM, Zoschke R, Schunemann D (2019) Ribosome-Associated Chloroplast SRP54 Enables Efficient Cotranslational Membrane Insertion of Key Photosynthetic Proteins. Plant Cell 31: 2734–2750

Ito K, Akiyama Y (2005) Cellular functions, mechanism of action, and regulation of FtsH protease. Annual Review of Microbiology 59: 211–231

Janska H, Kwasniak M, Szczepanowska J (2013) Protein quality control in organelles - AAA/FtsH story. Biochim Biophys Acta 1833: 381–387

Järvi S, Suorsa M, Paakkarinen V, Aro EM (2011) Optimized native gel systems for separation of thylakoid protein complexes: novel super- and mega-complexes. Biochem J 439: 207–214

Jarvis P, Lopez-Juez E (2013) Biogenesis and homeostasis of chloroplasts and other plastids. Nat Rev Mol Cell Biol 14: 787–802

Kato Y, Hyodo K, Sakamoto W (2018) The Photosystem II Repair Cycle Requires FtsH Turnover through the EngA GTPase. Plant Physiol 178: 596–611

Kato Y, Sun X, Zhang L, Sakamoto W (2012) Cooperative D1 degradation in the photosystem II repair mediated by chloroplastic proteases in Arabidopsis. Plant Physiol 159: 1428–1439

Kikuchi S, Asakura Y, Imai M, Nakahira Y, Kotani Y, Hashiguchi Y, Nakai Y, Takafuji K, Bedard J, Hirabayashi-Ishioka Y, Mori H, Shiina T, Nakai M (2018) A Ycf2-FtsHi Heteromeric AAA-ATPase Complex Is Required for Chloroplast Protein Import. Plant Cell 30: 2677–2703

Komenda J, Tichy M, Prasil O, Knoppova J, Kuvikova S, de Vries R, Nixon PJ (2007) The exposed N-terminal tail of the D1 subunit is required for rapid D1 degradation during photosystem II repair in Synechocystis sp PCC 6803. Plant Cell 19: 2839–2854

Koussevitzky S, Stanne TM, Peto CA, Giap T, Sjogren LL, Zhao Y, Clarke AK, Chory J (2007) An Arabidopsis thaliana virescent mutant reveals a role for ClpR1 in plastid development. Plant Mol Biol 63: 85–96

Langklotz S, Baumann U, Narberhaus F (2012) Structure and function of the bacterial AAA protease FtsH. Biochim Biophys Acta 1823: 40–48

Li Q, Wang X, Lei Y, Wang Y, Li B, Liu X, An L, Yu F, Qi Y (2022) Chloroplast envelope ATPase PGA1/AtFtsH12 is required for chloroplast protein accumulation and cytosol-chloroplast protein homeostasis in Arabidopsis. J Biol Chem 298: 102489

Lindahl M, Spetea C, Hundal T, Oppenheim AB, Adam Z, Andersson B (2000) The thylakoid FtsH protease plays a role in the light-induced turnover of the photosystem II D1 protein. Plant Cell 12: 419–431

Liu S, Zheng L, Jia J, Guo J, Zheng M, Zhao J, Shao J, Liu X, An L, Yu F, Qi Y (2019) Chloroplast Translation Elongation Factor EF-Tu/SVR11 Is Involved in var2-Mediated Leaf Variegation and Leaf Development in Arabidopsis. Front Plant Sci 10: 295

Liu X, Zheng M, Wang R, Wang R, An L, Rodermel SR, Yu F (2013) Genetic interactions reveal that specific defects of chloroplast translation are associated with the suppression of var2-mediated leaf variegation. J Integr Plant Biol 55: 979–993

Liu ZF, Yan HC, Wang KB, Kuang TY, Zhang JP, Gui LL, An XM, Chang WR (2004) Crystal structure of spinach major light-harvesting complex at 2.72 angstrom resolution. Nature 428: 287–292

Luciński R, Jackowski G (2013) AtFtsH heterocomplex-mediated degradation of apoproteins of the major light harvesting complex of photosystem II (LHCII) in response to stresses. J Plant Physiol 170: 1082–1089

Malnoё A, Wang F, Girard-Bascou J, Wollman FA, de Vitry C (2014) Thylakoid FtsH protease contributes to photosystem II and cytochrome b6f remodeling in Chlamydomonas reinhardtii under stress conditions. Plant Cell 26: 373–390

Mulo P, Pursiheimo S, Hou CX, Tyystjarvi T, Aro EM (2003) Multiple effects of antibiotics on chloroplast and nuclear gene expression. Functional Plant Biology 30: 1097–1103

Nishimura K, Kato Y, Sakamoto W (2017) Essentials of Proteolytic Machineries in Chloroplasts. Mol Plant 10: 4–19

Nixon PJ, Michoux F, Yu J, Boehm M, Komenda J (2010) Recent advances in understanding the assembly and repair of photosystem II. Ann Bot 106: 1–16

Ogura T, Tomoyasu T, Yuki T, Morimura S, Begg KJ, Donachie WD, Mori H, Niki H, Hiraga S (1991) Structure and function of the ftsH gene in Escherichia coli. Res Microbiol 142: 279–282

Okuno T, Ogura T (2013) FtsH protease-mediated regulation of various cellular functions. Subcell Biochem 66: 53–69

Okuno T, Yamanaka K, Ogura T (2006) An AAA protease FtsH can initiate proteolysis from internal sites of a model substrate, apo-flavodoxin. Genes to Cells 11: 261–268

Pipitone R, Eicke S, Pfister B, Glauser G, Falconet D, Uwizeye C, Pralon T, Zeeman SC, Kessler F, Demarsy E (2021) A multifaceted analysis reveals two distinct phases of chloroplast biogenesis during de-etiolation in Arabidopsis. Elife 10

Pribil M, Labs M, Leister D (2014) Structure and dynamics of thylakoids in land plants. Journal of Experimental Botany 65: 1955–1972

Puchades C, Rampello AJ, Shin M, Giuliano CJ, Wiseman RL, Glynn SE, Lander GC (2017) Structure of the mitochondrial inner membrane AAA+ protease YME1 gives insight into substrate processing. Science 358

Qi Y, Liu X, Liang S, Wang R, Li Y, Zhao J, Shao J, An L, Yu F (2016) A Putative Chloroplast Thylakoid Metalloprotease VIRESCENT3 Regulates Chloroplast Development in Arabidopsis thaliana. J Biol Chem 291: 3319–3332

Qi YF, Wang XM, Lei P, Li HM, Yan LR, Zhao J, Meng JJ, Shao JX, An LJ, Yu F, Liu XY (2020) The chloroplast metalloproteases VAR2 and EGY1 act synergistically to regulate chloroplast development in Arabidopsis. Journal of Biological Chemistry 295: 1036–1046

Reinbothe C, Bartsch S, Eggink LL, Hoober JK, Brusslan J, Andrade-Paz R, Monnet J, Reinbothe S (2006) A role for chlorophyllide a oxygenase in the regulated import and stabilization of light-harvesting chlorophyll a/b proteins. Proceedings of the National Academy of Sciences of the United States of America 103: 4777–4782

Rutschow H, Ytterberg AJ, Friso G, Nilsson R, van Wijk KJ (2008) Quantitative proteomics of a chloroplast SRP54 sorting mutant and its genetic interactions with CLPC1 in Arabidopsis. Plant Physiol 148: 156–175

Sakamoto W, Miura E, Kaji Y, Okuno T, Nishizono M, Ogura T (2004) Allelic characterization of the leaf-variegated mutation var2 identifies the conserved amino acid residues of FtsH that are important for ATP hydrolysis and proteolysis. Plant Molecular Biology 56: 705–716

Sakamoto W, Tamura T, Hanba-Tomita Y, Murata M (2002) The VAR1 locus of Arabidopsis encodes a chloroplastic FtsH and is responsible for leaf variegation in the mutant alleles. Genes to Cells 7: 769–780

Sakamoto W, Zaltsman A, Adam Z, Takahashi Y (2003) Coordinated regulation and complex formation of yellow variegated1 and yellow variegated2, chloroplastic FtsH metalloproteases involved in the repair cycle of photosystem II in Arabidopsis thylakoid membranes. Plant Cell 15: 2843–2855

Schünemann D (2004) Structure and function of the chloroplast signal recognition particle. Curr Genet 44: 295–304

Schünemann D, Gupta S, Persello-Cartieaux F, Klimyuk VI, Jones JDG, Nussaume L, Hoffman NE (1998) A novel signal recognition particle targets light-harvesting proteins to the thylakoid membranes. Proceedings of the National Academy of Sciences of the United States of America 95: 10312–10316

Silva P, Thompson E, Bailey S, Kruse O, Mullineaux CW, Robinson C, Mann NH, Nixon PJ (2003) FtsH is involved in the early stages of repair of photosystem II in Synechocystis sp PCC 6803. Plant Cell 15: 2152–2164

Sun JL, Li JY, Wang MJ, Song ZT, Liu JX (2021) Protein Quality Control in Plant Organelles: Current Progress and Future Perspectives. Mol Plant 14: 95–114

Suno R, Niwa H, Tsuchiya D, Zhang XD, Yoshida M, Morikawa K (2006) Structure of the whole cytosolic region of ATP-dependent protease FtsH. Molecular Cell 22: 575–585

Susek RE, Ausubel FM, Chory J (1993) Signal transduction mutants of Arabidopsis uncouple nuclear CAB and RBCS gene expression from chloroplast development. Cell 74: 787–799

Tomoyasu T, Gamer J, Bukau B, Kanemori M, Mori H, Rutman AJ, Oppenheim AB, Yura T, Yamanaka K, Niki H, Hiraga S, Ogura T (1995) Escherichia-Coli Ftsh Is a Membrane-Bound, Atp-Dependent Protease Which Degrades the Heat-Shock Transcription Factor Sigma(32). Embo Journal 14: 2551–2560

Walker D, Moshbahi K, Vankemmelbeke M, James R, Kleanthous C (2007) The role of electrostatics in colicin nuclease domain translocation into bacterial cells. Journal of Biological Chemistry 282: 31389–31397

Wang P, Grimm B (2015) Organization of chlorophyll biosynthesis and insertion of chlorophyll into the chlorophyll-binding proteins in chloroplasts. Photosynthesis Research 126: 189–202

Wang P, Liang FC, Wittmann D, Siegel A, Shan SO, Grimm B (2018) Chloroplast SRP43 acts as a chaperone for glutamyl-tRNA reductase, the rate-limiting enzyme in tetrapyrrole biosynthesis. Proceedings of the National Academy of Sciences of the United States of America 115: E3588–E3596

Wang R, Zhao J, Jia M, Xu N, Liang S, Shao J, Qi Y, Liu X, An L, Yu F (2018) Balance between Cytosolic and Chloroplast Translation Affects Leaf Variegation. Plant Physiol 176: 804–818

Wang X, Li Q, Zhang Y, Pan M, Wang R, Sun Y, An L, Liu X, Yu F, Qi Y (2022) VAR2/AtFtsH2 and EVR2/BCM1/CBD1 synergistically regulate the accumulation of PSII reaction centre D1 protein during de-etiolation in Arabidopsis. Plant Cell Environ 45: 2395–2409

Weber ER, Hanekamp T, Thorsness PE (1996) Biochemical and functional analysis of the YME1 gene product, an ATP and zinc-dependent mitochondrial protease from S-cerevisiae. Molecular Biology of the Cell 7: 307–317

Yu F, Liu X, Alsheikh M, Park S, Rodermel S (2008) Mutations in SUPPRESSOR OF VARIEGATION1, a factor required for normal chloroplast translation, suppress var2-mediated leaf variegation in Arabidopsis. Plant Cell 20: 1786–1804

Yu F, Park S, Rodermel SR (2004) The Arabidopsis FtsH metalloprotease gene family: interchangeability of subunits in chloroplast oligomeric complexes. Plant Journal 37: 864–876

Yu F, Park S, Rodermel SR (2005) Functional redundancy of AtFtsH metalloproteases in thylakoid membrane complexes. Plant Physiol 138: 1957–1966

Yuan J, Kight A, Goforth RL, Moore M, Peterson EC, Sakon J, Henry R (2002) ATP stimulates signal recognition particle (SRP)/FtsY-supported protein integration in chloroplasts. J Biol Chem 277: 32400–32404

Zaltsman A, Ori N, Adam Z (2005) Two types of FtsH protease subunits are required for chloroplast biogenesis and photosystem II repair in Arabidopsis. Plant Cell 17: 2782–2790

Zhang D, Kato Y, Zhang LG, Fujimoto M, Tsutsumi N, Sodmergen, Sakamoto W (2010) The FtsH Protease Heterocomplex in Arabidopsis: Dispensability of Type-B Protease Activity for Proper Chloroplast Development. Plant Cell 22: 3710–3725

Zheng M, Liu X, Liang S, Fu S, Qi Y, Zhao J, Shao J, An L, Yu F (2016) Chloroplast Translation Initiation Factors Regulate Leaf Variegation and Development. Plant Physiol 172: 1117–1130

Zhu D, Xiong HB, Wu JH, Zheng CH, Lu DD, Zhang LX, Xu XM (2022) Protein Targeting Into the Thylakoid Membrane Through Different Pathways. Frontiers in Physiology 12

Ziehe D, Dunschede B, Schunemann D (2018) Molecular mechanism of SRP-dependent light-harvesting protein transport to the thylakoid membrane in plants. Photosynth Res 138: 303–313

